# Intrinsic RNA targeting constrains the utility of CRISPR-Cas13 systems

**DOI:** 10.1101/2022.05.14.491940

**Authors:** Zexu Li, Zihan Li, Xiaolong Cheng, Xiaofeng Wang, Shixin Ma, Shengnan Wang, Zhiyan Lu, Han Zhang, Wenchang Zhao, Zhisong Chen, Yingjia Yao, Lumen Chao, Wei Li, Teng Fei

**Affiliations:** College of Life and Health Sciences, Northeastern University, Shenyang 110819, People’s Republic of China; Key Laboratory of Data Analytics and Optimization for Smart Industry (Northeastern University), Ministry of Education, Shenyang 110819, People’s Republic of China; Center for Genetic Medicine Research, Children’s National Hospital, 111 Michigan Ave NW, Washington, DC, 20010, USA; Department of Genomics and Precision Medicine, George Washington University, 111 Michigan Ave NW, Washington, DC, 20010, USA

**Keywords:** Cas13, CRISPR, RNA, lentivirus, screen

## Abstract

CRISPR-Cas13 systems have been adapted as versatile toolkits for RNA-related applications. Here we systematically evaluate the performance of several popular Cas13 family effectors (Cas13a, Cas13b and Cas13d) under lentiviral vectors and reveal surprisingly differential defects and characteristics of these systems. Using RNA immunoprecipitation sequencing, transcriptome profiling, biochemistry analysis, high-throughput CRISPR-Cas13 screening and machine learning approaches, we determine that each Cas13 system has its intrinsic RNA targets in mammalian cells. Viral process-related host genes can be targeted by Cas13 and affect production of fertile lentiviral particles, thereby restricting the utility of lentiviral Cas13 systems. Multiple RNase activities of Cas13 are involved in endogenous RNA targeting. Unlike target-induced nonspecific collateral effect, intrinsic RNA cleavage can be specific, target-independent and dynamically tuned by varied states of Cas13 nucleases. Our work provides guidance on appropriate use of lentiviral Cas13 systems and further raises cautions about intrinsic RNA targeting during Cas13-based basic and therapeutic applications.

## INTRODUCTION

The type VI CRISPR-Cas13 systems have been repurposed as powerful toolkits for RNA-targeting applications such as RNA knockdown, editing, imaging, detection and gene therapy (Abudayyeh et al., 2019; Abudayyeh et al., 2016; East-Seletsky et al., 2016; Gootenberg et al., 2017; Perculija et al., 2021; Powell et al., 2022; Terns, 2018; Yang et al., 2019). Well characterized and commonly used Cas13 family effectors include Cas13a, Cas13b and Cas13d (Abudayyeh et al., 2017; Cox et al., 2017; Konermann et al., 2018; Smargon et al., 2017; Yan et al., 2018). Recent studies on Cas13X/Y, Cas13bt and Cas7-11 further expanded such RNA-targeting tools (Kannan et al., 2022; Ozcan et al., 2021; Xu et al., 2021). Each Cas13 RNA nuclease achieves site-specific RNA targeting via a single guide RNA (sgRNA) consisting of a short cognate direct repeat (DR) sequence and a 20∼30 bp target-matching spacer sequence within the CRISPR RNA (crRNA) (O’Connell, 2019). Once activated by pairing between crRNA sequence and complementary single-stranded RNA (ssRNA) target, Cas13 effector exerts target RNA cleavage in *cis*. Meanwhile, activated Cas13 often exhibits a *trans* or collateral cleavage activity by which Cas13 can degrade surrounding non-target ssRNA. Such collateral effect may compromise the precision of Cas13-mediated RNA targeting and induce cellular toxicity, however, this property has been harnessed to amplify readout signals in RNA detection applications (Gootenberg *et al*., 2017; Myhrvold et al., 2018). Both *cis* and *trans* RNA cleavage activities of Cas13 are mediated by conserved higher eukaryotes and prokaryotes nucleotide-binding (HEPN) domain. Moreover, Cas13 proteins also possess an additional RNase activity that enables pre-crRNA processing and is enzymatically distinct from RNA-guided on-target *cis* or nonspecific *trans* activities (Abudayyeh *et al*., 2016; East-Seletsky *et al*., 2016; Liu et al., 2017; Zhang et al., 2019).

There are several ways of introducing these programmable CRISPR effectors into target cells for genetic manipulation or gene therapy. Among them, viral vector-based delivery represents one of the mainstream approaches (Doudna, 2020; Komor et al., 2017). Since lentivirus can infect almost any mammalian cell types, those widely used CRISPR plasmids are often constructed under lentiviral vector backbones to achieve maximum target cell range in many basic and translational studies. For human gene therapy, adeno-associated virus (AAV) delivery system is preferably employed due to its optimal safety profile. In contrast to DNA-targeting enzymes such as Cas9, there have been limited reports using lentivirus-based Cas13 nucleases for either single gene manipulation or high-throughput screening applications, despite the vast enthusiasm to exploit and apply these tools in various contexts. A few cases are primarily implemented with Cas13d but quite rarely with Cas13a/b under lentiviral vectors (Abbott et al., 2020; Li et al., 2021; Wessels et al., 2020; Zhang et al., 2021). Apart from considerations on knockdown efficacy and collateral effect among different Cas13 nucleases, it remains enigmatic that whether or which Cas13 effectors can be readily applicable under lentiviral vectors.

Here we systematically investigated the performance and characteristics of three major Cas13 family effectors (Cas13a/b/d) that were packaged and delivered via lentiviral vectors. We found surprisingly dramatic difference between these lentiviral Cas13 systems and provided practical guidance to utilize these tools properly. Mechanistically, we revealed that Cas13 family nucleases exhibited different levels of intrinsic RNA targeting in mammalian cells, including a cohort of viral process-related genes in lentivirus-packaging cells. Such previously uncharacterized RNA processing activity on the host transcriptome not only affects lentiviral success of Cas13 systems, but also raises cautions on Cas13 implementation in target cells during both basic and translational applications.

## RESULTS

### Lentiviral Defects of One-vector Cas13 Systems

To directly compare and evaluate their performance on RNA interference, we firstly cloned LwCas13a (referred to hereafter as Cas13a), PspCas13b (Cas13b) and RfxCas13d (Cas13d) into the same lentiviral vector backbone lentiCRISPR v2 (lentiv2) by replacing Cas9 and sgRNA cassette with corresponding Cas13 components (Figure 1A). Transient transfection of either nucleus-localized or cytoplasm-localized Cas13a/b/d all exhibited significant RNA interference as exemplified by two independent crRNAs targeting *EZH2* mRNA in HEK293FT cells (Figures 1B, S1A and S1B; Table S1). We then packaged lentiviruses with these vectors in HEK293FT cells and infected a broader range of cells that are refractory to transient transfection. Surprisingly, when we used puromycin to select Cas13-expressing clones after lentiviral infection in human lung adenocarcinoma A549 cells, Cas13d groups produced a significant amount of viable cells whereas both Cas13a and Cas13b groups had a scarcity of surviving cells left (Figure 1C). Similar phenomena were also observed when infecting multiple cell types such as breast cancer T47D cells and prostate cancer LNCaP cells using independent batches of lentiviruses (Figures S1C and S1D). The failure of gathering enough surviving cells for one-vector lentiviral Cas13a/b systems hindered their functional evaluation, although one-vector Cas13d system can be successfully delivered by lentiviral infection and the target transcripts were effectively knocked down in multiple cell types (Figures 1D and 1E).

**Figure 1.**
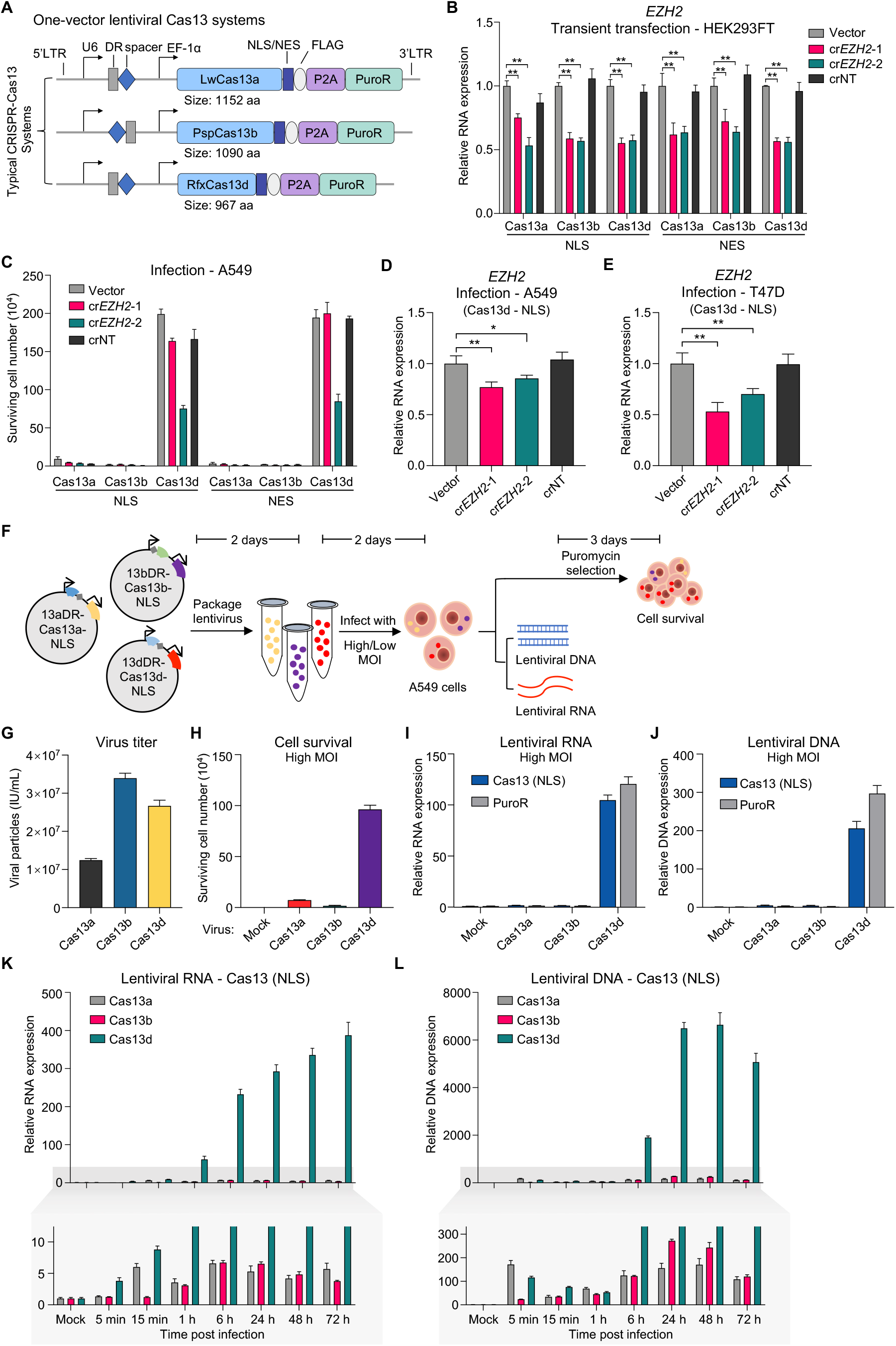
Lentiviral defects of one-vector Cas13 systems. (A) Schematic for one-vector lentiviral constructs expressing Cas13 effector and corresponding sgRNA cassette. (B) RNA knockdown activity of endogenous *EZH2* mRNA in HEK293FT cells by transient transfection of indicated one-vector lentiviral Cas13 plasmids, each with an empty vector, two independent crRNAs and a non-targeting (NT) crRNA. Values shown as mean ± SD with n = 3. \*\**p* < 0.01. (C) Surviving cell number after puromycin selection for A549 cells infected by indicated one-vector Cas13 lentiviruses, mean ± SEM with n = 3. (D-E) Knockdown activity of *EZH2* in A549 cells (D) or T47D cells (E) by one-vector Cas13d lentiviruses, mean ± SD with n = 3. \**p* < 0.05, \*\**p* < 0.01. (F) Schematic of evaluation assays for one-vector lentiviral Cas13 systems. (G) Virus titer of one-vector Cas13a/b/d lentiviruses, mean ± SEM with n = 3. (H) Surviving cell number against puromycin selection for A549 cells infected with indicated lentiviruses with high MOI, mean ± SEM with n = 3. Mock, uninfected cell group. (I) Lentiviral RNA expression of infected A549 cells by detecting NLS (the common tag for different Cas13 constructs) or PuroR elements, mean ± SD with n = 3. (J) Integrated lentiviral DNA levels from genomic DNA of infected A549 cells, mean ± SD with n = 3. (K-L) Time course examination of lentiviral RNA (K) or integrated DNA (L) post lentiviral infection in A549 cells, mean ± SD with n = 3. See also Figure S1 and Table S1.

To ascertain the lentiviral failure of one-vector Cas13a/b systems, we carefully dissected the whole infection processes to pinpoint the problematic steps (Figure 1F). Successful lentivector expression requires the following steps: (1) lentivirus packaging and collection; (2) cellular infection by co-incubation of virus with target cells; (3) lentiviral RNA reverse transcription into DNA and integration into host genome; and (4) expression of lentiviral gene cassette. We firstly quantified the viral particles packaged in HEK293FT cells and did not find dramatic difference between Cas13a/b/d systems (Figure 1G). Meanwhile, we did not observe apparent cellular toxicity in HEK293FT cells when expressing either Cas13 genes alone or together with packaging elements during lentivirus production (Figures S1E and S1F). After infection of A549 cells, we measured surviving cell number after puromycin selection, RNA expression of lentiviral elements (Cas13 or PuroR), and integrated lentiviral DNA elements (Cas13 or PuroR) from extracted genomic DNA. Consistent with initial observation, Cas13a/b infection did not produce viable puromycin-resistant cells no matter using high (>2) or low (0.3∼0.5) multiplicity of infection (MOI) of viruses, whereas Cas13d viruses were able to generate ample amount of surviving cells after selection (Figures 1H and S1G). Furthermore, in contrast to Cas13d, neither lentiviral RNA nor DNA was detected in Cas13a/b infected cells without puromycin selection (Figures 1I, 1J, S1H and S1I), indicating that Cas13a/b viruses are defective to integrate lentiviral DNA into the genome of target cells. Through a detailed time course analysis post infection, we observed a continuous accumulation of lentiviral RNA (initially from direct viral transfer and later from stable expression after genomic integration) and a significant plateau of corresponding DNA (resulting from reverse transcription and stable integration into the genome) for Cas13d (Figures 1K and 1L). However, Cas13a/b groups only exhibited a subtle increase in the early phase of infection and then dropped after around 48 hours for both RNA and DNA levels, which is likely a reflection of just lentiviral vector transfer and promiscuous reverse transcription but without successful genomic integration (Figures 1K and 1L). Moreover, we tested more lentiviral constructs with multiple crRNA inserts and found similar defects for Cas13a/b, suggesting a spacer-independent effect (Figures S1J-S1L). To rule out potential vector-specific effect, we employed another lentiviral vector backbone pHAGE-EF1α-puro to express Cas13a/b/d components and still observed similar phenomena (Figure S1M). These results revealed an unexpected lentiviral failure effect for one-vector Cas13a/b systems which may dramatically restrain their applications.

### Lentiviral Defects of Two-vector Lentiviral Cas13 Systems

We next constructed two-vector lentiviral Cas13 vectors by separately inserting Cas13a/b/d proteins and their corresponding sgRNA cassettes into lentiCRISPR v2 backbone in replacement of original Cas9 elements (named lenti-Cas13 and lenti-Cas13DR, respectively) (Figure 2A). Similar assays were performed to evaluate lentiviral infection for these Cas13 elements (Figure 2B). Interestingly, when lentivectors only expressed Cas13 protein without containing sgRNA cassette, Cas13b group produced a significant amount of viable cells after puromycin selection, which was in dramatic contrast to one-vector Cas13b system (Figures 2C and S2A). On the other hand, Cas13a- and Cas13d-expressing vectors showed similar results as their one-vector counterparts (Figures 2C and S2A). Lentiviral RNA and DNA quantification further confirmed that Cas13b/d-expressing vectors could execute fertile lentiviral infection while Cas13a-only vector still failed to integrate into target genome and express the transgenes (Figures 2D, 2E, S2B and S2C). In contrast, lentivectors expressing sgRNA cassette for Cas13a/b/d were all successfully delivered and expressed in target cells via lentiviral infection (Figures 2F and 2G), suggesting that Cas13 protein is the key for lentiviral failure effect.

**Figure 2.**
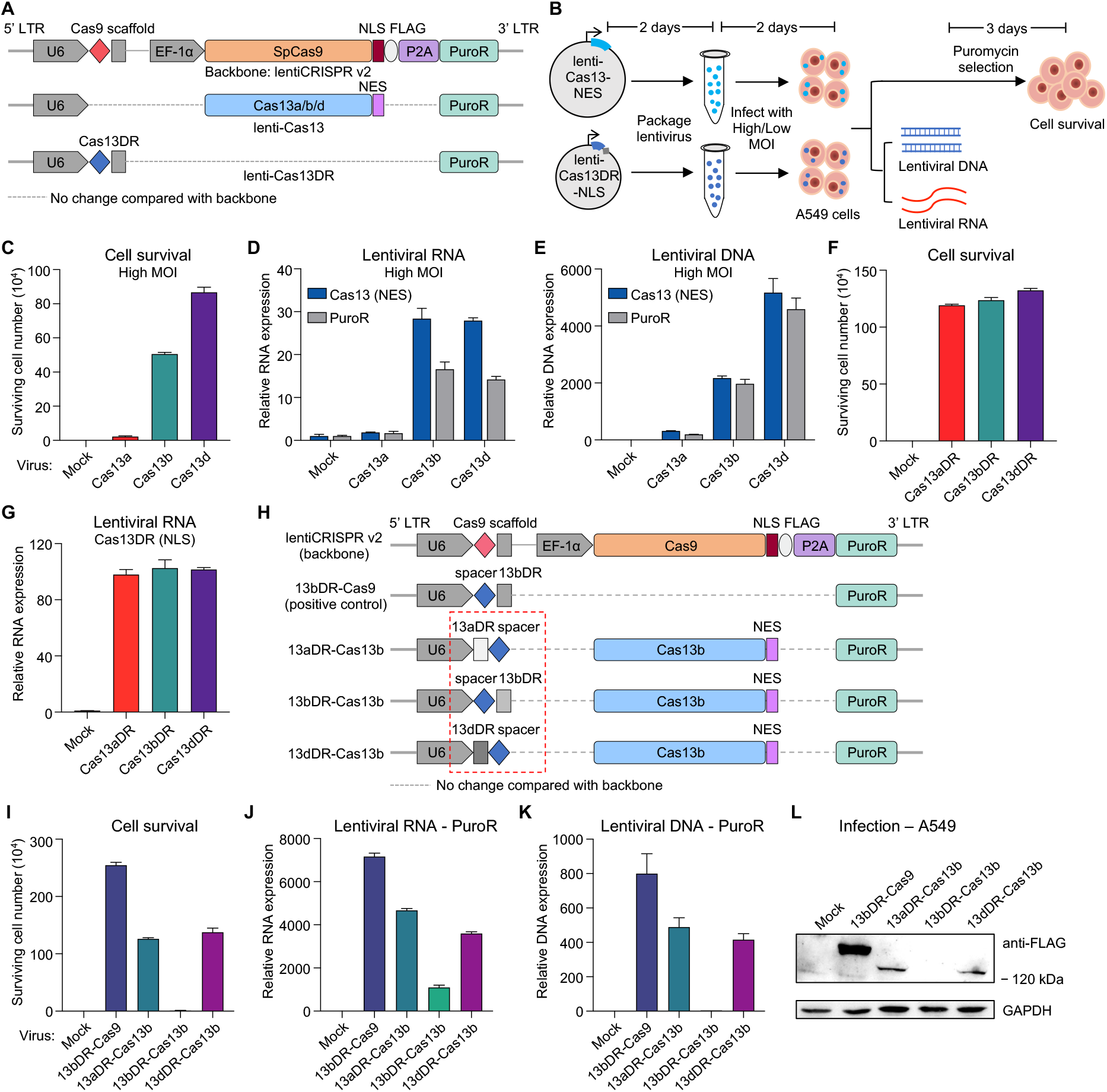
Lentiviral defects of two-vector Cas13 systems. (A) Schematic of two-vector lentiviral Cas13 constructs. (B) Schematic of evaluation assays for two-vector lentiviral Cas13 systems. (C-E) Assessment of surviving cell number (C), lentiviral RNA (D) and integrated lentiviral DNA (E) in A549 cells infected with Cas13-only lentiviruses at high MOI. (F-G) Surviving cell number (F) and lentiviral RNA level (G) after puromycin selection for A549 cells infected by Cas13DR lentiviruses of two-vector systems. (H) Schematic of one-vector lentiviral constructs containing Cas13b along with different Cas13DR. No spacer was inserted. (I-K) Assessment of surviving cell number (I), lentiviral RNA (J) and integrated lentiviral DNA (K) in A549 cells infected with indicated lentiviruses. (L) Western blot of protein extracts from A549 cells of untreated (Mock) or infected with indicated lentiviruses. See also Figure S2 and Table S1.

It is interesting that one-vector Cas13b displayed apparent lentiviral defect while two-vector Cas13b was lentivirally competent. Such discrepancy prompted us to hypothesize that the co-occurrence of Cas13b with its sgRNA cassette is critical to elicit lentiviral defect. To test this, we constructed a series of sgRNA cassette swapping vectors by changing the combination of Cas13b with different DR scaffolds (without specific spacer insert) in one-vector format (Figure 2H). Interestingly, lentiviral failure occurred only when Cas13b was conjugated with its own sgRNA cassette but not with other sgRNA architectures (Figures 2I-2L), suggesting that the binding of sgRNA scaffold to Cas13b protein is a requisite for such lentiviral defect. Using pHAGE-EF1α-puro as another independent lentivector backbone to express Cas13 proteins, we further confirmed the above findings on two-vector lentiviral Cas13 systems (Figures S2D and S2E).

Genomic integration of lentiviral vectors requires not only viral components but also host factors of infected cells to be recruited and functionalized. To explore whether lentiviral defect is due to virus itself or malfunctional host factors, we performed a co-infection experiment to see if lentivirally competent virus (sgRNA scaffold) could rescue the lentivector integration of deficient virus (Cas13a) in target cells (Figure S2F). Although viable cells post puromycin selection appeared for both Cas13a and Cas13b two-vector co-infection, lentiviral RNA expression of Cas13a was still very low and its sgRNA vector was responsible for puromycin resistance as evidenced by high lentiviral expression of Cas13aDR (NLS) element (Figures S2G-S2J). In contrast, both Cas13b and its sgRNA cassette were expressed in target cells (Figures S2G-S2J), indicating that two-vector Cas13b, but not one-vector version, is capable for lentiviral delivery. These data showed that competent virus co-infection in the same target cells could not rescue such lentiviral failure, suggesting that Cas13 virus itself but not host factors in target cells dictates the lentiviral defect. In addition, we also evaluated the protein stability of Cas13a/b/d in HEK293FT cells, and found that Cas13d protein was the most stable while Cas13a degraded very quickly (Figures S2K and S2L). Taken together, Cas13d is superior in both one-vector and two-vector formats for lentiviral delivery, and Cas13b is conditionally capable in only two-vector version while Cas13a is not usable for lentiviral system at all.

### Intrinsic RNA Targets of Different Cas13 Systems

Since Cas13 proteins are bona fide RNases and occasionally exhibit collateral RNA cleavage activity, we hypothesized that Cas13 also has intrinsic RNA targets in cells and these targets in lentivirus-packaging HEK293FT cells may contribute to lentiviral failure effect. To identify Cas13-associated RNA targets, we performed protein-RNA interaction analysis by RNA immunoprecipitation coupled with high-throughput sequencing (RIP-seq) in HEK293FT cells transiently expressing different Cas13 components with or without lentiviral defect (Figure S3A). Compared to Cas9 and empty vector control, all Cas13 groups possessed an increased number of RNA binding sites over the transcriptome (ranging from 1,973 to 4,121 strong peaks with stringent cutoff) (Figure S3B and Table S2, see Methods). A majority of Cas13-binding sites were located in exon and 3’-UTR regions of the transcript (Figure S3C). All Cas13 family displayed similar GA-enriched binding motifs which were quite distinct from that of Cas9-associated RNAs (Figure S3D). These Cas13-associated peaks converged on 1321 to 2492 genes for each Cas13 group and a few genes were shared between groups (Figures 3A and 3B). These genes were significantly enriched for fundamentally biological processes such as “translation” and “RNA processing” (Figure S3E). In addition, “viral gene expression” and “viral process” also appeared within the top 20 enriched terms (Figure S3E). Considering their relatedness to lentiviral defect, we elaborately examined the relevance between Cas13-associated genes and broader viral processes. Interestingly, we found that many viral process-related terms were enriched among Cas13-bound genes and such enrichment positively correlated with the potential of lentiviral defect (Figure 3C). Moreover, among the top consensus Cas13-associated RNA transcripts, we found a significant portion of viral process-related genes as reported previously (Besnard et al., 2016; Chen et al., 2016; Kumar et al., 2017; Rahim et al., 2018; Sadat et al., 2014; Saha et al., 2011; Sammaibashi et al., 2018; Santoni et al., 2020; Xia et al., 2019; Yoh et al., 2015) (Figure 3D). These data suggest that intrinsic targeting of those viral process-related gene transcripts in HEK293FT cells may affect the production of competent lentiviruses and subsequently lead to lentiviral failure in target cells.

**Figure 3.**
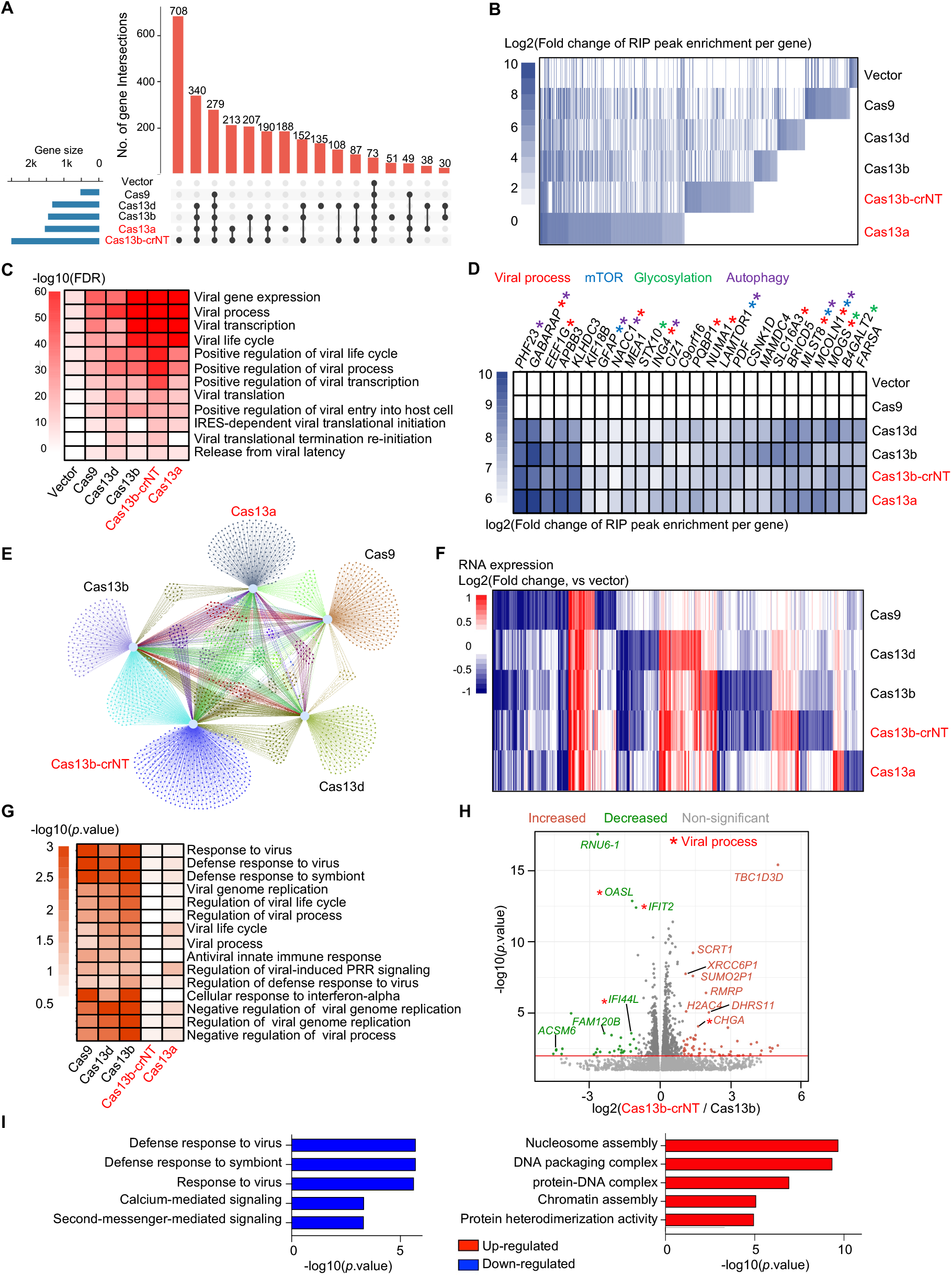
Intrinsic RNA targets of different Cas13 effectors. (A-B) Overlap (A) and heatmap (B) of genes associated with strong Cas13-bound peaks over the transcriptome. Vector: lentiv2-w/o Cas9. (C) Viral process-related gene ontology (GO) categories enriched among Cas13-associated peaks. (D) Top consensus Cas13-associated RNA gene transcripts with viral process-related genes denoted by red asterisk. (E-F) Overlap (E) and heatmap (F) of differentially expressed genes in Cas9-or Cas13-expressing HEK293FT cells. (G) Viral process-related GO categories enriched among Cas9- or Cas13-upregulated genes. (H-I) Volcano plot (H) and top 5 enriched GO categories (I) of down- and up-regulated differentially expressed genes, Cas13b-crNT versus Cas13b. See also Figure S3; Tables S1, S2 and S3.

We further performed transcriptome profiling by RNA-seq in those Cas9- and Cas13-expressing cells (Figure S3A). Compared to vector control, hundreds of genes were significantly changed (|fold change|>1.5, adjusted *p* value <0.05) upon Cas9 or Cas13 expression with certain concordance between Cas13 samples (Figures 3E, 3F and S3F; Table S3). Functional categories such as “nucleosome” or “response to virus” were significantly enriched among Cas13-regulated genes (Figures S3G and S3H). Interestingly, when specifically looking into viral process-related terms, we found a clear inverse correlation between term enrichment and lentiviral defect potential for up-regulated genes across tested samples (Figure 3G), suggesting that lentivirus-competent groups might up-regulate a cohort of host genes to boost lentiviral capacity while lentivirus-deficient samples fail to do so. Considering the adaptive immunity and anti-virus nature of CRISPR-Cas systems, we posited that such host gene regulation by Cas proteins might be reflective of feedback response during evolutionary interaction between host milieu, co-evolved virus and anti-virus machinery. To further pinpoint candidate genes that could potentially mediate lentiviral defect, we directly compared samples between lentivirus-deficient Cas13b-crNT (one-vector format) and lentivirus-competent Cas13b (protein-only part in two-vector version) that closely resembled in components but clearly differed in lentiviral capacity (Figures 3H and S3I). Several top differential genes were known to be implicated in viral regulation (Figure 3H). Furthermore, virus-related terms were among the top five significantly enriched functional categories in under-represented genes of Cas13b-crNT sample compared to Cas13b group (Figure 3I), indicating that those down-regulated virus-relevant genes undergoing Cas13-mediated RNA cleavage may underlie the lentiviral defect of Cas13b-crNT. Despite that both Cas13-associated and -regulated RNA targets were enriched for virus-related genes, we did not observe significant correlation between Cas13 binding and Cas13-induced gene expression change (data not shown). It is probably because that enzyme-substrate interaction during Cas13-mediated intrinsic RNA cleavage might obey a classic “hit-and-run” model in which Cas13 only transiently interacts with its cleaved intrinsic RNA substrates, thereby making it difficult to capture strong Cas13 RIP signals for those targets. Furthermore, in addition to direct RNA cleavage, the effects of Cas13 binding to its targeted RNA may also be executed through non-enzymatic mechanisms such as RNA localization and protein translation control. Taken together, these Cas13-associated and -regulated intrinsic RNA targets might be impactful on biological properties of target cells, including lentiviral capacity for lentivirus-packaging cells.

### Pre-crRNA Cleavage Activity Underlies the Lentiviral Defect of Cas13a

Cas13 proteins possess two major types of RNase activities: HEPN domain-dependent *cis*/*trans* RNA cleavage activity and pre-crRNA processing activity. Using LwCas13a, we tried to determine whether such RNase activity is responsible for lentiviral failure effect. We firstly pinpointed key amino acid residues in LwCas13a that are necessary for *cis*/*trans* or pre-crRNA cleavage using prior knowledge or by homology assessment with other reported Cas13a proteins (Figure S4A) (Abudayyeh *et al*., 2016; East-Seletsky *et al*., 2016; Liu *et al*., 2017). We then generated several point mutations of LwCas13a that ablated either HEPN (abbreviated as Cas13a-H), pre-crRNA processing (Cas13a-R) or both (Cas13a-HR) activities (Figures 4A and 4B). *In vitro cis*/*trans* and pre-crRNA cleavage assays using purified Cas13 proteins further confirmed the function of these mutants (Figures S4B-S4F). We then packaged these Cas13a variants into lentiviruses in HEK293FT cells and infected A549 cells to examine their lentiviral capacity. Despite similar viral titers, wild-type (WT) lentiviral Cas13a was unable to yield puromycin-resistant cells and concordantly showed very low lentiviral RNA expression and DNA integration (Figures 4C-4F). Interestingly, Cas13a-R and Cas13a-HR mutant could fully rescue the lentiviral failure phenotype, while Cas13-H mutant was partially effective in reversing lentiviral defect only in high MOI condition (Figures 4C-4F). These results suggest that the pre-crRNA processing activity is the pivotal source for the promiscuously intrinsic RNA targeting in HEK293FT cells that underlies lentiviral defect of Cas13a.

**Figure 4.**
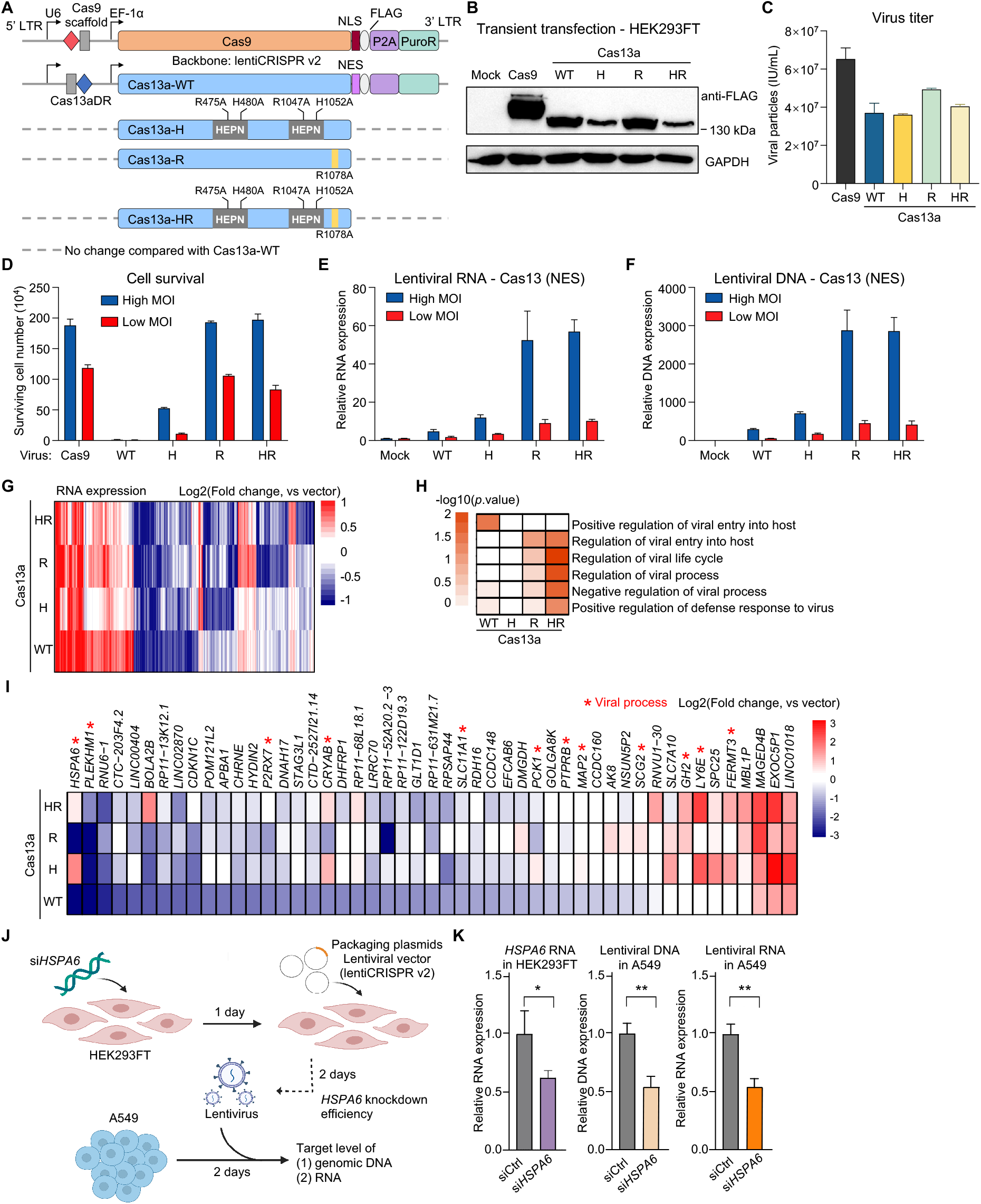
Pre-crRNA cleavage activity underlies lentiviral defect of Cas13a. (A) Schematic of one-vector lentiviral Cas13a-WT and different RNase-deficient variant constructs. No spacer was inserted. (B) Western blot of protein extracts from HEK293FT cells transiently transfected with indicated constructs. (C) Virus titer of indicated lentiviruses, mean ± SEM with n = 3. (D-F) Assessment of surviving cell number (D), lentiviral RNA (E) and integrated lentiviral DNA (F) in A549 cells infected with indicated lentiviruses at high or low MOI. (G) Heatmap of differentially expressed genes in Cas13a variant-expressing samples. (H) Viral process-related GO categories enriched among up-regulated genes by Cas13a variants. (I) Heatmap of representative up-regulated or de-repressed genes in RNase-deficient Cas13a variants compared with Cas13a-WT. Viral process-related genes are marked by red asterisk. (J) Schematic for evaluating lentiviral capacity of *HSPA6* knockdown by siRNA in HEK293FT cells. Lentiviruses produced in either control (siCtrl) or *HSPA6* knockdown HEK293FT cells were used to infect A549 cells and lentiviral RNA or DNA level was determined. (K) Assessment of *HSPA6* knockdown efficiency in HEK293FT cells (left), lentiviral DNA (center) and lentiviral RNA level (right) in A549 cells by measuring a region spanning NLS and FLAG tag in lentiCRISPR v2 vector, mean ± SD with n = 3. See also Figure S4, Tables S1 and S3.

To explore potential RNA targets affected by different RNase activities, we performed RNA-seq in HEK293FT cells transiently expressing these Cas13a variants (Table S3). Each variant affected RNA expression of hundreds of genes with some common hits as well as some variant-specific targets (Figures 4G and S4G). Varied functional terms were enriched in up-or down-regulated genes for different variants (Figures S4H and S4I). Consistent with the lentiviral phenotype, viral process-related terms were primarily enriched in up-regulated genes for lentivirally competent Cas13a-R and -HR samples but less for lentivirally compromised Cas13-WT or -H groups (Figure 4H), suggesting that RNase-deficient Cas13a mutants might de-repress those virus-related genes to reverse the lentiviral defect caused by Cas13a-WT. Among those top de-repressed genes in Cas13a mutants versus Cas13a-WT, several genes were reported to participate in viral regulation (Figure 4I). For example, *HSPA6*, encoding a heat shock protein of Hsp70 family, can be induced upon enterovirus A71 infection and is required for viral life cycle (Su et al., 2021). Here we showed that *HSPA6* was significantly repressed by Cas13a-WT but dramatically de-repressed in Cas13a-H and -HR mutants (Figure 4I), indicating that endogenous *HSPA6* might undergo HEPN-dependent RNA cleavage by Cas13a. Interestingly, *HSPA6* was significantly induced upon lentivector transfection in HEK293FT cells but got significantly repressed when introducing different Cas13 proteins via lentivectors, whereas its close family members *HSPA1* and *HSPA8* did not display such lentivector- and Cas13-dependent effects (Figure S4J). Moreover, we found that *HSPA6* knockdown by small interfering RNA (siRNA) in HEK293FT cells significantly attenuated lentiviral capacity as evidenced by reduced lentiviral RNA and DNA levels in target A549 cells (Figures 4J and 4K). Notably, *HSPA6* was a HEPN-dependent pan-Cas13 target that could generally affect lentiviral capacity of all the Cas13 effectors tested. On the other hand, the virus-related Cas13a targets among de-repressed genes in Cas13a-R or -HR mutants might collectively underlie specific lentiviral defect of Cas13a-WT. Overall, these data suggest that Cas13-mediated endogenous RNA cleavage on viral process-related gene transcripts in packaging cells is crucial for lentiviral capacity of Cas13 tools.

### Endogenous RNA Cleavage by Cas13a

To provide direct evidence that Cas13 can target intrinsic RNA for cleavage without target-matching crRNA, we established an *in vitro* RNA cleavage assay by co-incubation of purified Cas13a variant proteins solely with target RNA fragments of interest. As hit-and-run model indicated a potentially transient interaction between Cas13 and its cleaved intrinsic RNA targets, we selected several gene transcripts with differential Cas13a binding strength for the test (Figure 5A). In addition, we also included *HSPA6* which we have validated as a HEPN-dependent functional Cas13 target (Figures 4I-4K), although its Cas13 binding was barely detected in RIP-seq (Figure 5A). Some of them displayed up-regulated or de-repressed expression pattern in RNase-deficient Cas13a mutants versus Cas13-WT according to RNA-seq while some genes such as *PRL13A* and *VPS11* did not despite strong Cas13 binding (Figure 5B). Potential targetable RNA fragments were chosen primarily according to RIP-seq tracks (denoted by red rectangle in Figure 5A; Table S1), obtained via *in vitro* transcription, and further co-incubated with purified Cas13 proteins for *in vitro* RNA cleavage. Using *EGFP* RNA fragment as an irrelevant control, we did not observe apparent RNA degradation by either Cas9 or Cas13a-WT (Figure 5C). Meanwhile, neither we saw clear Cas13a-mediated RNA cleavage for tested RNA fragments of *PRL13A* and *VPS11* (Figure 5C), consistent with their steady expression levels under Cas13a variants (Figure 5B). These results indicate that strong physical association with Cas13 does not necessarily result in RNA cleavage, consistent with the hit-an-run model. Interestingly, Cas13a-WT caused significant degradation of tested RNA fragments from *LY6E*, *PYCR3*, *DHRSX* and *HSPA6*, whereas such cleavage was greatly blunted by Cas13a-HR and to a lesser extent by Cas13a-H or -R mutants (Figure 5D), suggesting that both HEPN and pre-crRNA processing activities could be involved in such intrinsic RNA cleavage. Although these cleaved RNA generally exhibited concordant upregulation or de-repression in cells expressing Cas13a-HR, the cleavage strength *in vitro* by Cas13 variants was not always consistent with expression change *in vivo*. It is probably because that the choice of tested RNA fragment greatly affects substrate selection by different RNase activity and endogenous RNA structure differs to that of purified RNA. We hypothesized that differential Cas13 conformation in varied activation state might affect such intrinsic RNA cleavage. Indeed, using *in vitro* assay, we found that *DHRSX* RNA cleavage by Cas13a-WT was attenuated in Cas13a:crRNA sample (lane 5 vs. lane 4) but greatly enhanced upon Cas13a:crRNA:on-target RNA ternary complex formation (lane 6) in which on-target *cis* and bystander *trans* cleavage activities were fully unleashed (Figure S5). Interestingly, in contrast to on-target RNA cleavage, intrinsic RNA cleavage by Cas13a did not exhibit clear band pattern which resembled that from *trans* activity (Figures 5D and S5). These data suggest that conformational change of Cas13 elicited by formation of binary Cas13:crRNA or ternary Cas13:crRNA:on-target RNA complex may affect substrate selection and thus alter RNA cleavage pattern as well as kinetics for those crRNA-independent endogenous RNA cleavage.

**Figure 5.**
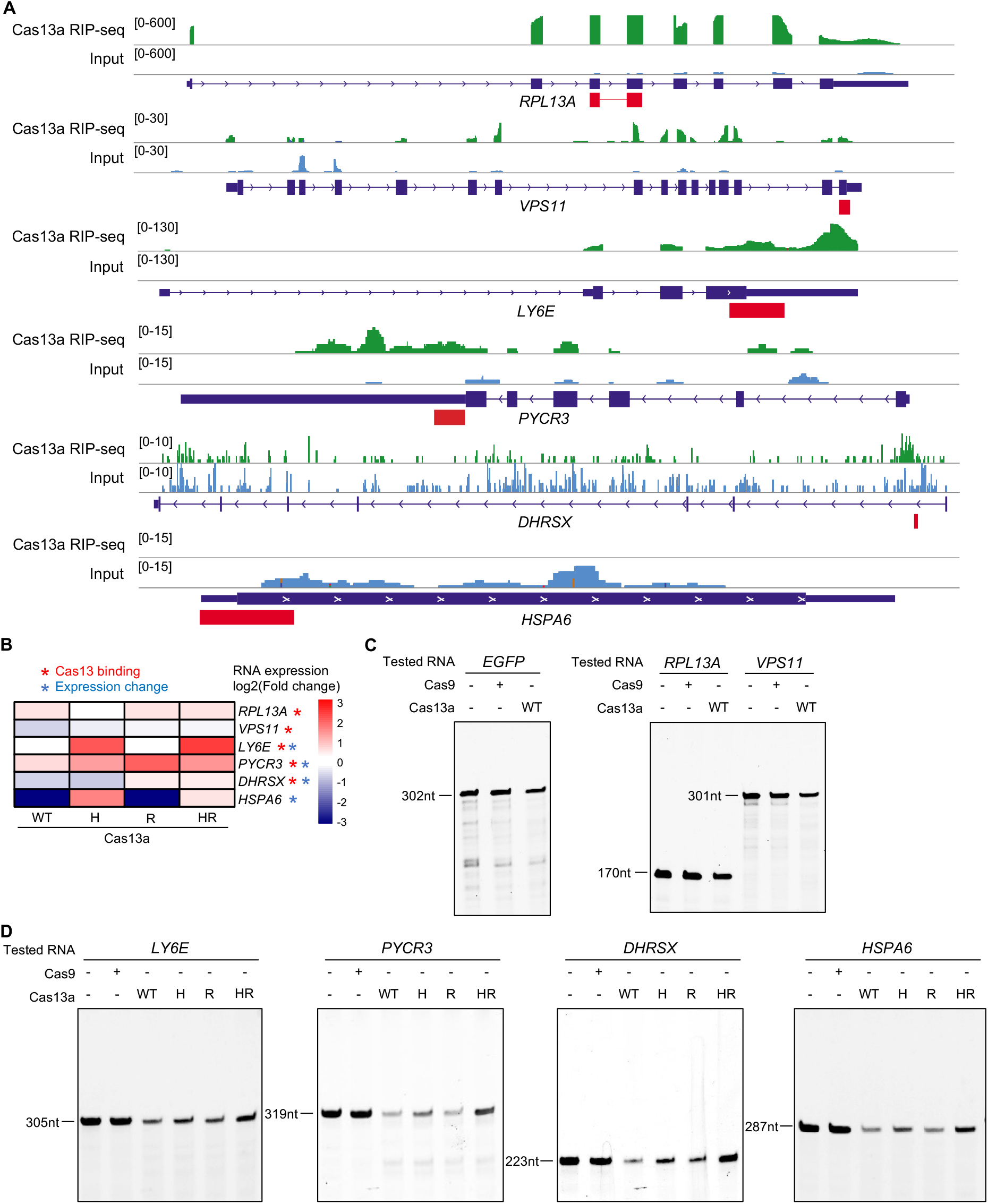
Endogenous RNA cleavage by Cas13a. (A) RIP-seq tracks of indicated genes for Cas13a. Red rectangle denotes RNA fragment used for *in vitro* cleavage assay. (B) Heatmap showing RNA expression pattern of indicated genes. (C) *In vitro* RNA cleavage for indicated RNA fragments by Cas13a-WT or Cas9. *EGFP* serves as an irrelevant control. (D) *In vitro* RNA cleavage assay for indicated RNA fragments by Cas13a variants. See also Figure S5; Tables S1, S2 and S3.

### Evaluation of Cas13 Performance by High-throughput Screening Approach

Competent lentivector is a pre-requisite for most of the high-throughput CRISPR screens. Based on our above findings, Cas13a in either one-vector or two-vector format and one-vector Cas13b were incapable for lentiviral Cas13 screens. Thus, we proceeded to evaluate the performance of Cas13d (one-vector or two-vector) and Cas13b (two-vector) during high-throughput screening application (Figure 6A). We selected 192 protein-coding genes and 122 cancer-related long noncoding RNAs (lncRNAs) as target transcripts (Figure S6A), and designed two independent sgRNA libraries for either Cas13d or Cas13b system with 10,830 and 11,936 crRNAs, respectively (Figure S6B; Table S4). Lentiviruses delivering these sgRNAs in either one-vector or two-vector format were produced in HEK293FT cells, and screening was performed in human melanoma A375 cells in duplicate to identify functional hits affecting cell fitness (Figure 6B; Table S5, see Methods). For two-vector systems, lentiviral sgRNA libraries were also introduced into wild-type (WT) Cas13-null cells as another level of control. We calculated log2 fold-change (lfc) of crRNA abundance of indicated time point compared to that of plasmid pool. Interestingly, in contrast to two-vector Cas13d samples, one-vector Cas13d groups displayed a dramatic dispersion of crRNA distribution even for the virus library and samples of early time point (Day 5) before functional selection in cells (Figures 6C and S6C). Considering the lentiviral defect for Cas13a/b systems, we posited that the deviated crRNA distribution might also be a reflection of lentiviral failure for specific subset of crRNA-containing one-vector Cas13d viruses. Such defect and distorted crRNA representation may further impede the analysis of true biological effect of interrogated genes in the screen. Indeed, when focusing on core essential genes during fitness screen, we observed significantly negative selection (selection end-Day 33 vs. selection start-Day 5) in two-vector Cas13d samples but not in two-vector Cas13d-null or one-vector Cas13d groups (Figures 6D and 6E), as illustrated by β score distribution computed by MAGeCK-VISPR algorithm (Li et al., 2015). These data indicate that only two-vector Cas13d system was capable to identify correct genes during fitness screen. Top selected protein-coding genes and lncRNA genes were highlighted with several previously unknown cell fitness regulators (Figures 6F and 6G). Facilitated by these two-vector Cas13d screening data, we further designed a machine learning model named DeepCas13 to predict Cas13d-based on-target and off-target effect of given crRNAs (Cheng et al., 2021; manuscript under consideration).

**Figure 6.**
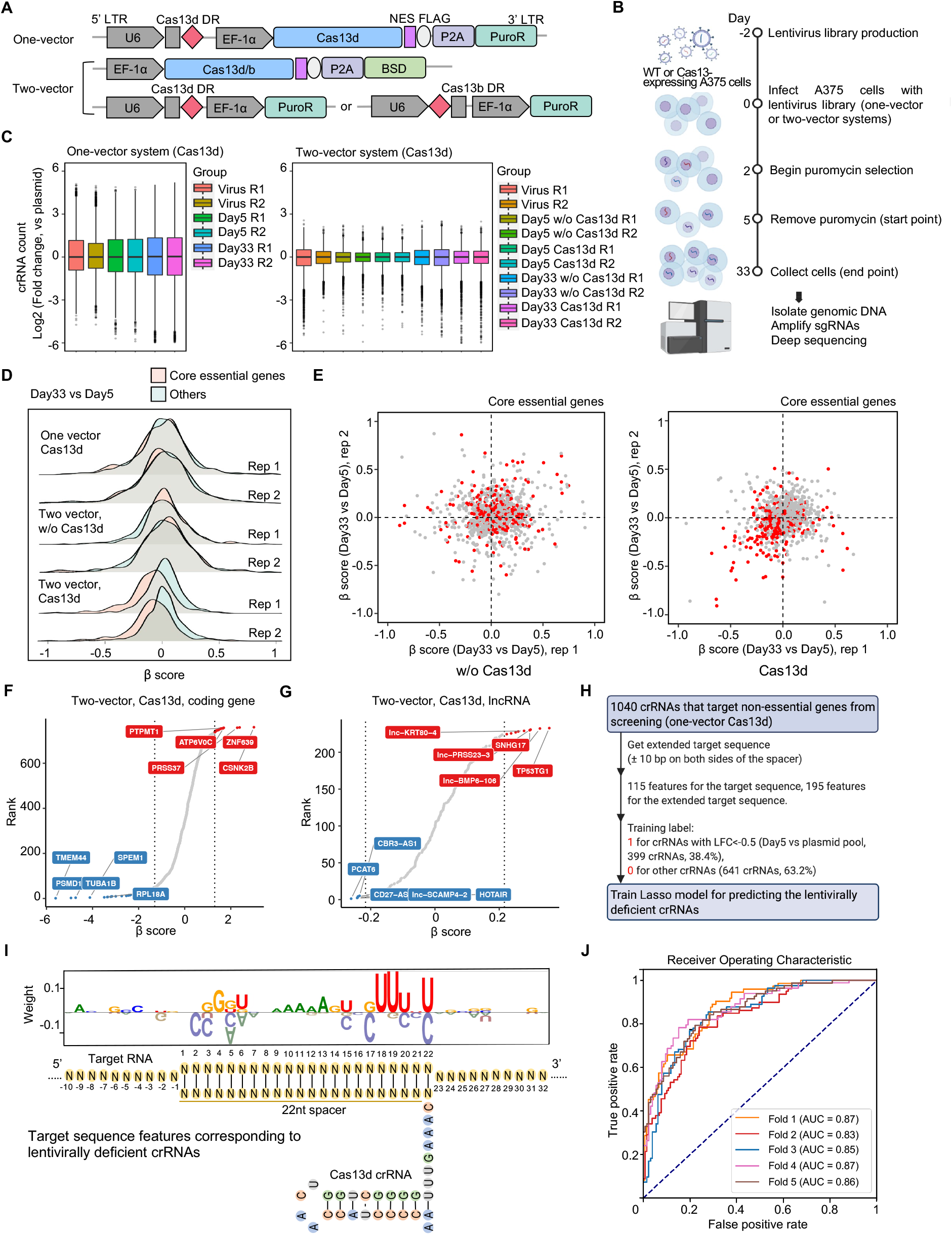
Evaluation of Cas13 performance during high-throughput screening. (A) Schematic of lentiviral vectors in one-vector or two-vector format for CRISPR-Cas13 screens. (B) Schematic of procedures for Cas13-based cell fitness screens. (C) The distribution of crRNA abundance across samples of different time points compared to plasmid pool for one-vector or two-vector Cas13d screen. (D) The β score distribution of core essential genes versus other genes at early (Day 5) or end point (Day 33) of the screens. (E) Scatter plot of β score change for target transcripts between two biological replicates in Cas13d-null or Cas13d-expressing conditions of two-vector Cas13d screens. Red dots indicate core essential genes. (F) Rank-ordered presentation of targeted protein-coding genes for their essentiality according to β score. The top selected genes were highlighted. (G) Rank-ordered presentation of targeted lncRNA genes for their essentiality according to β score. The top selected lncRNAs were highlighted. (H) Schematic of lasso-based machine learning model to determine crRNA-specific features underlying lentiviral defect. (I) A logo for highly represented sequence features embedded in early depleted crRNAs. (J) Receiver Operator Characteristic (ROC) curves showing the predictive power of lasso model using 5-fold cross validation. See also Figure S6; Table S4 and S5.

For two-vector Cas13b screens, although there was not apparent crRNA count distortion in virus pool and early screening stages (Figure S6D), we still failed to observe significant shift toward negative selection for crRNAs targeting core essential genes in either Cas13b-expressing or Cas13b-null samples (Figures S6E and S6F). The less efficacy of two-vector Cas13b screens than that of Cas13d may be reflective of weaker RNA interference capability of Cas13b *per se* according to previous studies (Konermann *et al*., 2018; Wessels *et al*., 2020), or imperfect selection of crRNAs due to lack of optimized gRNA design rules for Cas13b. Collectively, these results suggest that two-vector lentiviral Cas13d system is the most suitable for massively parallel sgRNA analysis in high-throughput screening application.

### Lentiviral Defect Elicited by Specific crRNAs in One-vector Cas13d System

Although empty vector of one-vector Cas13d showed no evident lentiviral defect (Figure 1), we did observe disproportional representation of some crRNAs among samples before cellular selection in one-vector Cas13d screens (Figures 6C and S6G), again implying the existence of somewhat crRNA-specific lentiviral defect. We postulated that the crRNAs that failed to be properly encapsulated into virus or integrated into the genome due to lentiviral defect should be enriched among those early depleted crRNA population when the true biological selection had not been fully initiated. To investigate the rules of crRNA-dependent lentiviral defect, we developed a lasso-based machine learning model and analyzed the enriched sequence features of extended target region (± 10 bp on both sides of the spacer) for those crRNAs that target non-essential genes but rather displayed strong depletion in the early stage (Day 5 vs. plasmid pool) of one-vector Cas13d screens (Figure 6H, see Methods). Interestingly, we found significant sequence features within the spacer region that might predict the potential crRNA-specific lentiviral failure of one-vector Cas13d system (Figure 6I). Using 5-fold cross validation, we confirmed that this lasso-based scoring model successfully predicted most of the early loss of matched crRNAs for non-essential genes in the screen (average AUC score= 0.86) (Figure 6J). To avoid potential lentiviral defect during crRNA design, we also implemented this lasso model into our DeepCas13 web server at http://deepcas13.weililab.org/ by calculating a lentiviral score to predict the potential of lentiviral defect for any given Cas13d crRNA. These results suggest that certain sequence features within crRNAs may additionally alter the binding kinetics of Cas13d with its substrate and elicit differential patterns of intrinsic RNA targeting that lead to compromised lentiviral capability. Such effects should be taken into consideration for one-vector Cas13d crRNA design.

## DISCUSSION

Programmable RNA targeting CRISPR-Cas13 systems greatly expand the dimensions of gene editing tools at the RNA level, and hold great potential in advancing both basic research and human gene therapy. Here we systematically evaluated the performance and property of three major types of Cas13 family effectors (Cas13a/b/d) under lentiviral vectors, and provided a practical guidance to utilize these tools appropriately. More importantly, we found that the lentiviral defect of Cas13 systems result from previously unrecognized intrinsic host RNA targeting by Cas13 effectors. Such endogenous RNA targeting activity not only constrains the utility of lentiviral Cas13 systems, but may also lead to other unexpected effects depending on specific RNA targets and target cell types. These findings call for in-depth consideration and additional evaluation for Cas13-based tools especially when using them in human clinics.

Lentiviral vectors enable efficient gene transfer and expression in almost any cell types. Unlike other CRISPR-based tools, studies using lentiviral Cas13 systems are comparatively limited, which is previously unexplained and significantly impedes the progression of Cas13 field. By systematical evaluation, we found that different Cas13 systems exhibit differential levels of lentiviral defect. Cas13a protein itself is sufficient to cause harmful intrinsic RNA targeting relevant to viral processes, thereby hindering the production of competent lentivirus. Cas13b induces lentiviral defect as long as it complexes with its cognate sgRNA scaffold regardless of spacer sequence. Interestingly, Cas13d generally does not exhibit lentiviral defect in either Cas13-DR coexistence one-vector format or separated two-vector system, however, Cas13d can also be lentivirally compromised when certain crRNA pairing with specific RNA targets with special sequence features in virus-packaging HEK293FT cells. The subtle conformational change of Cas13:crRNA:on-target RNA ternary complex elicited by specific crRNA:target pairing may expose the catalytic surface of Cas13d to endogenous RNA targets in host cells, which underlies the lentiviral defect. Taken together, for lentiviral usage of Cas13 systems, we recommend the following rules: (1) avoid using LwCas13a; (2) apply PspCas13b only in two-vector format but some crRNA:target pairing may not work; 3) cautiously use one-vector RfxCas13d provided careful crRNA design and target region selection to avoid special sequence features; and (4) preferentially use RfxCas13d in two-vector format. In addition, Cas13d has a generally superior knockdown efficiency compared to Cas13b or Cas13a as revealed by single gene assay and high-throughput screening evaluation by our study and other reports (Konermann *et al*., 2018; Wessels *et al*., 2020).

We further explored the molecular mechanism for the observed lentiviral defect and revealed a previously unappreciated but prevalent effect of endogenous RNA targeting by Cas13 nucleases. As a bona fide RNA binding protein, Cas13 can associate with a plethora of endogenous RNA transcripts and potentially affect their expression by several mechanisms such as RNA cleavage, stability, subcellular localization and RNA translation control. Classic *cis* and *trans* RNA cleavage require crRNA:target pairing which then leads to Cas13 activation by exposing catalytic pocket in HEPN domains to on-target or bystander RNA. Moreover, Cas13 can also cleave long pre-crRNA transcript to produce mature crRNA without formation of Cas13:crRNA:on-target RNA ternary activation complex (Abudayyeh *et al*., 2016; East-Seletsky *et al*., 2016; Konermann *et al*., 2018). Here we surprisingly observed that Cas13 by itself can selectively cleave endogenous RNA transcripts even in the absence of crRNA or crRNA:target pairing-mediated Cas13 activation, indicating that different Cas13 states (protein along, Cas13:crRNA binary complex, or Cas13:crRNA:on-target RNA ternary complex) possess differential capability and selectivity for intrinsic RNA targeting. Mutagenesis of key residues in HEPN domains or residues important for pre-crRNA processing compromised the cleavage strength of endogenous RNA, suggesting the implication of these known RNase activities for innate RNA targeting. Notably, we still cannot rule out other uncharacterized RNase activities that may be involved in this process since double domain mutation of known RNase activities (HR) did not always fully abrogate such endogenous RNA cleavage (Figure 5D). Therefore, even for the applications with catalytically dead Cas13 (dCas13), researchers should also be aware of potential unwanted effect derived from intrinsic RNA targeting. On the other hand, loss-of-function mutation for pre-crRNA processing seems to fully rescue the lentiviral defect of Cas13a although HEPN domain mutation also has partial effect, indicating that a subset of intrinsic targets responsible for lentiviral production are mainly regulated through pre-crRNA processing activity of Cas13a.

Complete activation of Cas13 by perfect crRNA:target pairing often induces a collateral effect that leads to nonspecific cleavage of bystander ssRNA. Such collateral effect was originally observed in bacterial cells for Cas13a (Abudayyeh *et al*., 2016), but was recently reported for more Cas13 effectors including Cas13d (Ai et al., 2022; Kelley et al., 2021; Li et al., 2022; Shi et al., 2021; Tong et al., 2021; Wang et al., 2019b). Compared with collateral cleavage, intrinsic RNA targeting we described here is fundamentally distinct in the following ways: (1) intrinsic RNA targeting is more prevalent as long as Cas13 protein is present, whereas collateral effect requires the presence and abundance of fully activated Cas13:crRNA:on-target RNA ternary complex; (2) more RNase domains and activities of Cas13 may be involved in intrinsic RNA targeting, while collateral RNA cleavage is exclusively mediated by the two HEPN domains of Cas13; (3) the cleavage effect of intrinsic RNA targeting is comparatively subtle and selective; whereas collateral RNA cleavage is drastic and nonspecific; (4) the influence of intrinsic RNA targeting is not limited to RNA cutting but may extend to broader RNA metabolism processes such as RNA translation or localization control, whereas collateral effect is only restricted to RNA cleavage; and (5) the phenotypic effect of intrinsic RNA targeting can be latent (e.g., lentiviral defect) or unspecified depending on given transcriptome milieu and target cell types, whereas collateral effect often causes apparent cellular toxicity (Figure S6H).

The presence of collateral effect significantly impedes the application of Cas13 nucleases for RNA knockdown purpose. Here our findings on intrinsic RNA targeting raise another layer of cautions that should be considered when applying Cas13 effectors in both basic and translational contexts. In addition to lentiviral defect, other unexpected influence from intrinsic RNA targeting may also exist depending on Cas13 activation status, transcriptome profile and target cell types. Thus, it is advisable to comprehensively evaluate the effect of intrinsic RNA targeting especially in Cas13-based therapeutic scenarios.

There are several limitations of our current study. First, we preferentially focused on the most popular Cas13a/b/d systems to date in our study although several novel RNA targeting CRISPR tools emerged recently such as Cas13X/Y, Cas13bt and Cas7/11 (Kannan *et al*., 2022; Ozcan *et al*., 2021; Xu *et al*., 2021). Moreover, we primarily utilized HEK293FT cells and lentiviral defect to characterize the targets and functional effects of intrinsic RNA targeting by Cas13. Further investigations are needed for more RNA-targeting tools in more diverse cell types to unveil broader phenotypic effects from intrinsic RNA targeting. Second, the detailed molecular mechanism remains unclear on how intrinsic RNA targeting is affected by differential activation states of Cas13. Structural analysis of multiple Cas13 complexes will help to resolve this issue. Third, despite guidance on how to use lentiviral Cas13 systems and machine learning models to predict lentivirally defective crRNAs for one-vector Cas13d, we have not provided a definite solution to perfectly remove the intrinsic RNA targeting activity of Cas13. With better understanding of the molecular mechanism, rational mutagenesis or directed protein evolution will be needed to remove such unwanted intrinsic RNA targeting and meanwhile keep precise on-target RNA processing capacity of Cas13 effectors.

## MATERIALS AND METHODS

### Plasmid construction

The following plasmid backbones were used in this study:

lentiCRISPR v2 (lentiv2, Addgene #52961): a lentiviral vector backbone, U6 promoter expresses Cas9 sgRNA scaffold; EF1α promoter expresses Cas9 and PuroR (puromycin resistance) with P2A linker;

lentiGuide-puro (Addgene #52963): an empty lentiviral vector expresses Cas9 sgRNA under U6 promoter with puromycin resistance;

lenti-Cas9-blast (Addgene #52962): a lentiviral vector expresses Cas9 protein and blasticidin resistance under EF1α promoter.

pHAGE-EF1α-puro: an empty lentiviral vector backbone, EF1α promoter drives expression of inserted cDNA; SV40 promoter expresses PuroR;

pcDNA3.1: a derivative of pcDNA3.1-HA (Addgene #128034) vector, without HA tag; lentiv2-w/o Cas9: a derivative of the lentiCRISPR v2 vector with Cas9 open reading frame (ORF) removal;

pET-28a(+): bacterial vector for expression of N-terminally 6xHis-tagged proteins with a thrombin site.

The coding regions of LwCas13a (*Leptotrichia wadeii* Cas13a) (Gootenberg *et al*., 2017), PspCas13b (*Prevotella sp. P5-125* Cas13b) (Cox *et al*., 2017) and RfxCas13d (*Ruminococcus flavefaciens* XPD3002 Cas13d) (Konermann *et al*., 2018) were inserted into lentiCRISPR v2 (replacing Cas9 ORF) and pHAGE-EF1α-puro lentiviral expression vector between XbaI and BamHI restriction enzyme sites with C-terminal fused FLAG tag and nucleoplasmin NLS or HIV NES signal. Cas13 DR scaffold was cloned by replacing Cas9 sgRNA cassette in lentiCRISPR v2 backbone. For pHAGE-EF1α-puro vector, the fragments of U6 promoter along with Cas13 DR cassette were inserted in SpeI restriction enzyme site. For two-vector screening plasmids, Cas13 sgRNA cassette was inserted into lentiGuide-puro to replace original Cas9 sgRNA cassette. Cas13 expression cassette was inserted into lenti-Cas9-blast to replace Cas9 ORF. With such strategy, the following plasmids were constructed:

lenti-9DR-Cas13a-NES, lenti-9DR-Cas13b-NES, lenti-9DR-Cas13d-NES; lenti-13aDR-Cas9-NLS, lenti-13bDR-Cas9-NLS, lenti-13dDR-Cas9-NLS; lenti-13aDR-Cas13a-NLS/NES, lenti-13bDR-Cas13b-NLS/NES, lenti-13dDR-Cas13d-NLS/NES; lenti-13aDR-Cas13b-NES, lenti-13dDR-Cas13b-NES; pHAGE-Cas13a-NLS, pHAGE-Cas13b-NLS, pHAGE-Cas13d-NLS; pHAGE-13aDR-Cas13a-NLS, pHAGE-13bDR-Cas13b-NLS, pHAGE-13dDR-Cas13d-NLS, lenti-Cas13b-NES-blast, lenti-Cas13d-NES-blast, lentiGuide-13bDR-puro, lentiGuide-13dDR-puro. 9DR denotes Cas9 sgRNA cassette and 13DR denotes Cas13 sgRNA cassette.

The following Cas13a variant plasmids were generated by introducing the following point mutations:

lenti-13aDR-Cas13a (WT)-NES;

lenti-13aDR-Cas13a (H)-NES, point mutations: R475A /H480A /R1047A /H1052A; lenti-13aDR-Cas13a (R)-NES, point mutation: R1078A;

lenti-13aDR-Cas13a (HR)-NES, point mutations: R475A /H480A /R1047A /H1052A /R1078A.

For protein purification, Cas9 and Cas13a variants were cloned into prokaryotic expression vector pET-28a(+) to generate the following plasmids:

pET-28a-Cas9; pET-28a-Cas13a (WT); pET-28a-Cas13a (H); pET-28a-Cas13a (R); and pET-28a-Cas13a (HR).

### Cell culture

HEK293FT, A375 (human melanoma cell line), T47D (human breast cancer cell line), LNCaP (human prostate cancer cell line) and A549 (human lung cancer cell line) cells were obtained from American Type Culture Collection (ATCC). All cells were regularly tested negative for mycoplasma contamination and maintained in DMEM (SEVEN Biotech) supplemented with 10% fetal bovine serum (ExCell) and 1% penicillin– streptomycin (Solarbio). Cells were passaged every 2-4 days to maintain exponential growth and were kept in a humidity-controlled incubator at 37°C with 5% CO_2_.

### Transient transfection in cells

HEK293FT cells were plated at a density of 200,000 cells per well in 12-well plate and transfected at > 50% confluency with 1 µg of expression plasmids using Neofect^TM^ DNA transfection reagent according to the manufacturer’s protocol. Transfected cells were harvested 48-72 hour (h) post-transfection for downstream analysis such as western blot or gene expression analysis.

### Lentivirus production and infection

DNA Transfection was performed in 6-well plates by Lipofectamine^TM^ 2000 Transfection Reagent with a mix of 1.5 μg lentiviral plasmid, 0.75 μg pCMVR8.74, and 0.5 μg pMD2.G. Two days after transfection, medium was harvested and centrifuged at 3000 rpm for 5 min to remove cell debris. Aliquot and store the virus supernatant at −80°C before use. Lentivirus titration was determined by qPCR using the Lentivirus Titer Kit (Applied Biological Materials, Abm) following the manufacturer’s instruction. The titer of virus can be calculated from online lentiviral titer calculator at http://www.abmgood.com/High-Titer-Lentivirus-Calculation.html. To test Cas13-based lentivirus infection in mammalian cells, A549, T47D or LNCaP cells were seeded at a density of 200,000 cells in 12-well plate, and infected at >30% confluency with lentivirus at a high multiplicity of infection (MOI) (>2) or low MOI (0.3∼0.5). After incubation for 48 h, cells were harvested for DNA and RNA extraction, respectively. Meanwhile, the medium was replaced with fresh culture medium containing appropriate concentration of puromycin and continue to culture for another 72 h prior to cell counting.

### Quantification of antibiotic-resistant cells, lentiviral RNA and DNA

For infection of A549, T47D and LNCaP cells, the selection concentration of puromycin was 1 µg/mL, 1.5 µg/mL and 3.5 µg/mL, respectively. The surviving cell number was counted using a hematocytometer. For quantification of lentiviral RNA and DNA, cells were harvested 48 h post infection for RNA and DNA extraction. *RPS28* served as internal control for both RNA and DNA quantification. Fold-change was calculated relative to *RPS28* control using **ΔΔ**Ct method. The primer set of qRT-PCR was designed on the common sequence (such as Cas13 adjacent NLS or NES tags, or on PuroR gene) across different samples. In cases of infection with different lentiviruses, appropriate primer combinations were used to assess their expression at RNA and DNA levels (Table S1).

### Protein expression and purification

Prokaryotic pET28a(+) plasmids expressing Cas9 or Cas13 proteins were transformed into *Escherichia coli*. Rosseta^TM^ 2(DE3)pLysS competent cells. After transformation, cells were plated on Luria-Bertani (LB) agar plate containing kanamycin and chloramphenicol, and incubated for 12∼16 h at 37°C. Pick up one colony from the plate, inoculate in 1L of LB media supplemented with antibiotics and shake at 37°C with 300 rpm. Until OD600 reached 0.4∼0.6, isopropyl β-D-thiogalactoside (IPTG) were added to a final concentration of 0.5 mM to induce protein expression for 14-16 h at 300 rpm in a pre-chilled 20°C shaker. After induction, the cells were harvested by centrifugation (5000 g, 4°C, 10 min) for later purification and stored at −80°C.

The whole process of protein purification was performed at 4°C. Briefly, cell pellet was resuspended in 15 mL of lysis buffer (20 mM Tris-HCl pH 8.0, 500 mM NaCl, 1 mM DTT, 5% glycerol, 1 mM PMSF, 1 mg/mL lysozyme), and lysed by sonication. The sonicated sample was then centrifuged at 14000 rpm for 10 min at 4°C and the supernatant was mixed with an equal volume of equilibration/wash buffer (50 mM sodium phosphate, 300 mM sodium chloride, 10 mM imidazole pH 7.4). HisPur^TM^ Cobalt Resin (Thermo Fisher, #89964) was utilized to pull-down the protein because the recombinant Cas protein contains His-tag. Washed protein extract was mixed with the prepared cobalt resin on end-over-end rotator for 30 min at 4°C followed by washing of the protein-bound cobalt resin twice in equilibration/wash buffer, and eluted by elution buffer (50 mM sodium phosphate, 300 mM sodium chloride, 150 mM imidazole pH 7.4). The eluted proteins were transferred into Zeba^TM^ Spin Desalting Columns (Thermo Fisher, #89890) for desalination, and ∼2 mL protein could be collected after centrifuging at 850 g for 2 min. Mix the protein with 10 mL Storage Buffer (600 mM NaCl, 50 mM Tris-HCl pH 7.5, 5% glycerol, 2 mM DTT), and transfer the mixture into Amicon Ultra-15 Centrifugal Filter Devices (Millipore, #UFC905008) to concentrate the protein and exchange the storage buffer as well. The purified proteins were quantified by BCA protein assay kit (Meilunbio, #MA0082) and SDS-PAGE followed by Coomassie Blue staining. Purified protein was stored at −80°C as 20 μL aliquots at a concentration of 2 mg/mL.

### Nucleic acid preparation

For preparation of RNA templates to test *cis* activity, *in vitro* transcription (IVT) was performed with T7 promoter-inclusive DNA templates. Briefly, *in vitro* transcription was performed with 4 μL 10x Transcription Buffer, 3.125 mM rNTPs, 50U T7 RNA Polymerase (Lucigen), 1 μL RNase Inhibitor and 1 µg DNA template in a 40 μL reaction system. After thorough mixing, the reaction system was incubated at 37°C for 1∼2 h. For preparation of crRNA and pre-crRNA, the DNA oligonucleotide containing reverse complementary sequence of crRNA was annealed to an oligo with T7 promoter sequence to form a partial duplex DNA as template of IVT for RNA production. For preparation of endogenous RNA targets to test cleavage, PCR amplifications were performed with synthesized DNA templates (Synbio Technologies) followed by IVT. The RNA products of IVT were purified by RNA Clean & Concentrator-5 Kit (Zymo research), and quantified by the high sensitivity RNA Qubit fluorometer (Thermo Fisher). All sequences used for *in vitro* cleavage are listed in Table S1.

### Cas13 cleavage assays

For testing *cis* activity of four LwCas13a variants, *in vitro* nuclease assays were performed with 100 nM purified Cas13a, 200 nM crRNA and 55 nM ssRNA templates in 1x CutSmart Buffer at 37°C for 1 h. For pre-crRNA processing reaction, the assay systems consisted of 200 nM purified Cas13a, 1x CutSmart Buffer and 290 nM pre-crRNA and reaction was performed at 37°C for 2 h. The reactions were stopped and resulting RNA was denatured with 2x RNA loading buffer (90% formamide, 0.5% EDTA, 0.1% xylene cyanol and 0.1% bromphenol blue) at 75°C for 5 min prior to gel electrophoresis. The samples were then loaded onto a 20% urea denaturing polyacrylamide gel with TBE buffer. For *trans* activity assay, the reaction system contained 50 nM purified Cas13a, 50 nM crRNA, 250 nM ssRNA fluorescence reporter, 1x CutSmart Buffer and 10 ng on-target RNA template. Fluorescence signal was dynamically measured at 37°C for 1 h by QuantStudio^TM^ 5 Real-Time PCR System (Thermo Fisher). Background-subtracted signals for each monitoring points were further normalized by subtraction of its initial value to make comparison between different conditions (arbitrary unit, a.u.) for the analysis. Visual detection was accomplished by imaging the tubes through E-Gel^TM^ Safe Imager^TM^ Real-Time Transilluminator (Thermo Fisher). *In vitro* endogenous RNA cleavage assays were performed with 200 nM purified Cas13a (or 200 nM purified Cas9 as control) and 200 ng endogenous RNA target fragment in 1x CutSmart Buffer. Reactions were incubated at 37°C for 2 h, then mixed with 2x RNA loading buffer at 75°C for 5 min for denaturation. Samples were analyzed on an 8% urea denaturing polyacrylamide gel with TBE buffer at 110 V for 50 min at 50°C. All the gels were stained with SYBR Gold (Invitrogen) for 10 min prior to imaging via Universal Hood II (BIO-RAD).

### Quantitative PCR (qPCR) analysis

To determine relative gene expression in RNA level, cells were harvested 48 h post transient transfection or infection and total RNA was extracted using UNIQ-10 Column Trizol Total RNA Isolation kit (Sangon). 1 µg of total RNA was then reverse-transcribed (RT) using random primers and MultiScribe Reverse Transcriptase (Thermo Fisher) at 25°C for 10 min, 37°C for 90 min, and 85°C for 5 min followed by qPCR using 2X UltraSYBR Mixture (CWBIO). House-keeping gene *RPS28* served as an internal control. Ct values were read out on a QuantStudio^TM^ 5 Real-Time PCR System (Thermo Fisher) with following program: 10 min 95°C for pre-denaturation, 40 cycles of 10 sec 95°C denaturation, 30 sec 60°C annealing and 30 sec 72°C elongation, with three 25 μL technical replicates in 96-well format. Fold-change was calculated relative to *RPS28* control using **ΔΔ**Ct method. For testing relative DNA levels, cells were harvested 48 h post infection and genomic DNA was extracted by phenol-chloroform and recovered by alcohol precipitation. Similar to the system and procedure mentioned in RT-qPCR, 100 ng genomic DNA was used instead as template in a 25 µL qPCR reaction mix.

### Western blot

For protein extraction, harvested cells were lysed in ice-cold RIPA lysis buffer (Beyotime) supplemented with protease inhibitor (Roche) at 4°C for 30 min. Then protein supernatant was collected by centrifuging at 14,000 rpm for 10 min at 4°C and mixed with 5X loading buffer. After boiling at 95°C for 5 min, the samples were analyzed by SDS-PAGE and blotted with indicated antibodies.

### Protein stability analysis

To compare protein stability between different Cas13 proteins *in vivo*, cycloheximide (CHX, a protein biosynthesis inhibitor) chase assays were performed with following protocol. 250,000 HEK293FT cells were plated in 6-well plate and transiently transfected at > 50% confluency with equal mole number of lenti-Cas13 expression plasmids using Neofect^TM^ DNA transfection reagent according to the manufacturer’s protocol. Each group of cells were divided into 5 equal parts in 24-well plate 24 h post transient transfection. After another 12 h culture, the cells were treated for 0, 2, 4, 6 and 8 h with 10 μg/mL CHX and collected to extract proteins. Western blot was used to examine the protein level change, and Image J software was used for gray analysis of images.

### RNA immunoprecipitation coupled with high-throughput sequencing (RIP-seq)

HEK293FT cells were plated in 15 cm plates and transfected at > 50% confluency with 25 µg of indicated plasmids using Neofect^TM^ DNA transfection reagent according to the manufacturer’s protocol. After 3 days transfection, cells were harvested by cross-linking with 0.3% formaldehyde for 10 min at room temperature and lysed with RIPA lysis buffer (50 mM Tris-HCl pH 7.6, 150 mM NaCl, 1 mM EDTA, 0.1% SDS, 1% NP-40, 0.5% sodium deoxycholate, protease inhibitor and RNase inhibitor) for 10 min on ice before sonication. The supernatant was collected and incubated with IgG (Santa Cruz #sc-2025) or FLAG antibodies (Invitrogen #MA1-91878) pre-incubated with Protein G beads (Invitrogen #10004D) at 4°C overnight. After washing with RIPA buffer (50 mM Tris-HCl pH 7.6, 1 M NaCl, 1 mM EDTA, 0.1% SDS, 1% NP-40 and 0.5% sodium deoxycholate), RNA was eluted from the beads with elution buffer (0.1 M NaHCO_3_ and 1% SDS, with proteinase K and RNase inhibitor) at room temperature for 10 min with gentle vortex. The eluted material was then de-cross-linked at 65°C for 45∼60 min in the presence of RNase inhibitor and proteinase K. Afterwards, RNA was purified using TRIzol LS reagent (Life Technologies) and treated with DNase I to remove any residual DNA. RIP RNA was used for library preparation. The libraries were constructed with TruSeq Stranded mRNA LT Sample Prep Kit (Illumina) according to the manufacturer’s instructions and then sequenced on Illumina PE150 sequencing platform (Novogene, China).

### Oligonucleotide library design for CRISPR screens

Due to intrinsic difference of targeting rules for Cas13b and Cas13d, separate oligo libraries targeting different regions of the same gene/transcript set were designed for either Cas13b screen or Cas13d screen. For Cas13d screens, one-vector or two-vector system utilized the same oligo library but different vector backbones. To evaluate the efficacy of Cas13-mediated RNA knockdown using cell fitness screens, we primarily chose known core essential genes / non-essential genes, cancer-related protein coding genes and potentially functional lncRNAs that are aberrantly expressed in cancers as target transcripts for oligo library design. Briefly, we selected 94 known “core essential genes” and 10 known “non-essential” genes whose perturbation are known to have strong (or no) effect on cell proliferation or viability from published resources (Hart et al., 2014; Wang et al., 2019a). In addition, we also included 88 other protein-coding genes with various functions in cancer, including oncogenes (e.g., *PIK3CA*, *PAK2* and *MYC*) or tumor suppressor genes (e.g., *RB1* and *CDKN1B*). For lncRNA genes, we selected lncRNAs that are overexpressed in multiple cancer types within the The Cancer Genome Atlas (TCGA) cohort (BRCA and PRAD). The lncRNA expression of each patient sample is downloaded from TCGA and a lncRNA is selected if it meets the following criteria: (1) the average log2 FPKM was greater than 2 for the 15% samples with the highest expression of this lncRNA; (2) log2 fold change (compared with normal samples) > 0.2; and (3) the expressions (FPKM) are greater than 10 in the corresponding cancer cell lines (T47D or LNCaP cells). In addition, 25 literature-curated lncRNAs with known functions in multiple cancers are also included (Du et al., 2013). To design crRNA, Ensembl gene and lncRNA annotations (version: GRCh38) are used to extract the sequences of the corresponding gene or lncRNA. For genes (or lncRNAs) with multiple transcripts, use transcripts with the corresponding RefSeq ID. We begin by enumerating all possible guides (22 bp for Cas13d or 30 bp for Cas13b) across the entire transcript, then remove guides that (1) map to more than one location in the human genome and transcriptome (allowing up to 1 mismatch) or (2) contain the BsmBI digestion site (“CGTCTC” or “GAGACG”). The remaining guides are randomly selected if they hit the greatest number of transcripts (“common” guides) in the same gene or lncRNA. If all the guides that hit the greatest number of transcripts are selected, the remaining guides that hit the second greatest number of transcripts will be randomly selected, and so on. For essential and non-essential genes, 35 guides are designed per gene. For other genes or lncRNAs, 15 guides are designed per gene or lncRNA. In total, 10,830 crRNAs were designed for Cas13d library and 11,936 crRNAs were included in Cas13b library. These oligos with flanking sequences were synthesized in a pooled format.

### CRISPR Library Synthesis and Construction

The pooled synthesized oligos (Synbio Technologies, China) were PCR amplified and then cloned into lentiGuide-13b/13dDR-puro or lenti-13dDR-Cas13d-NES via BsmBI site by Gibson Assembly. The ligated Gibson Assembly mix was transformed into self-prepared electrocompetent Stable *E. coli* cells (New England Biolabs) by electro-transformation to reach the efficiency with at least 100X coverage representation of each clone in the designed library. The transformed bacteria were then cultured in liquid LB medium for 16∼20 h at 30°C. The library plasmids were then extracted with EndoFree Maxi-prep Plasmid Kit (TIANGEN # 4992194).

### Pooled Genome-wide Cas13 Screens for Cell Fitness Gene Transcripts

Plasmid Cas13 libraries were firstly transfected along with pCMV8.74 and pMD2.G packaging plasmids into HEK293FT cells using Lipofectamine™ 2000 Transfection Reagent (Invitrogen #11668019) to generate pooled lentiviruses. Harvest virus-containing media at 72 h post transfection, and spin down the media at 1000 g for 5 min to remove the floating cells and cell debris. Carefully collect the virus supernatant, aliquot and store them at −80°C. Determine the virus titer and test MOI before proceeding to the screen. For two-vector Cas13b/d screens, Cas13b/d-expressing A375 cells were prepared in advance by infection of Cas13b/d-only lentiviruses (lenti-Cas13b/d-NES-blast) followed by blasticidin selection. Wild-type or Cas13-expressing cells (∼3×10^7^) were infected with one-vector Cas13d library or two-vector Cas13b/d sgRNA libraries with a low MOI (∼0.3) (Day 0). Two days later, select the infected cells with puromycin (1 µg/mL for A375 cells) for three days to get rid of non-infected cells (Day 5, start point of screen). The resulting cells were cultured in normal media for cell fitness screen until Day 33 (end point of screen). At least ∼300X coverage of cells were collected for samples of indicated time points and stored at −80°C before genomic DNA isolation and high-throughput sequencing for determining crRNA abundance.

### Genomic DNA isolation and sequencing

Wash cells with PBS and spin at 1000 rpm for 5 min. Discard supernatant. For around 5×10^6^ cells (∼500X coverage), add 1 mL lysis buffer (300 mM NaCl, 0.2% SDS, 2 mM EDTA, 10 mM Tris-HCl pH 8.0) in 2mL conical tube and 10 μL RNase A (10 mg/mL) to each sample followed by mixing and incubation at 65°C for 1 h. Then add 10 μL proteinase K (10mg/ml) to continue incubation at 55°C overnight (or 6 h). Mix with 1 mL phenol/chloroform/isoamyl alcohol solution (25:24:1) and centrifuge at 12000 rpm for 10 min before taking the upper phase. The genomic DNA was precipitated by isopropanol and washed by 75% ethanol before dissolved in nuclease-free water. To construct sequencing library, two rounds of PCR were performed. For the first round of PCR, perform 20-25 separate 100 µL reactions with 6-8 µg genomic DNA (maximum 10 µg per reaction) in each reaction using Q5 High-Fidelity DNA Polymerase (New England Biolabs) with the following primers: 1stF: 5’-AATGGACTATCATATGCTTACCGTAACTTGAAAGTATTTCG-3’; 1stR: 5’-GGAGTTCAGACGTGTGCTCTTCCGATCTCCAGTACACGACATCACTTTCCCAGT TTAC-3’. Combine the resulting amplicons together before proceeding to the next step. A second round of PCR aims to attach Illumina adaptors and to barcode samples. Perform the second round of PCR in a 100 µL reaction volume using 1 µL of the product from the first round of PCR for 10-12 cycles with the following primers: 2ndF: 5’-AATGATACGGCGACCACCGAGATCTACACTCTTTCCCTACACGACGCTCTTCCGATCTATCTTGTGGAAAGGACGAAACACC-3’; Index_R: 5’-CAAGCAGAAGACGGCATACGAGATNNNNNNNNGTGACTGGAGTTCAGACGTGT GCTCTTCCGATCT-3’ (N(8) are the specific index sequences). These PCR products were gel purified and pooled for high-throughput sequencing to determine crRNA abundance on Illumina PE150 sequencing platform (Novogene, China).

### Pooled Cas13 CRISPR screen analysis

The abundance of Cas13 gRNAs or crRNAs was calculated using the MAGeCK count module for the raw fastq files. The MAGeCK test module was applied to identify robust guide and gene-level enrichment with default parameters. The Robust Rank Aggregation (RRA) algorithm was also applied to normalize and rank the read counts. These gRNAs that target non-essential genes were used as control guides to estimate the size factor for normalization.

### RNA-seq and enrichment analysis

HEK293FT cells were transiently transfected with indicated constructs. After 72 hours, cells were harvested for RNA extraction and library preparation for stranded RNA-seq. Samples were sequenced on MGI DNBSEQ-T7 PE150 sequencing platform (Novogene, China). To quantify gene expression, pre-trimmed reads were aligned to the hg38 human reference genome with the UCSC known gene transcript annotation using HISAT2. Sequencing read coverage per gene was counted using HTSeq. Differentially expressed genes (DEGs) were identified using DESeq2. Genes with more than 1.5 fold change, Benjamini-Hochberg adjusted *p*-value < 0.05 and mean read count ≥ 5 were deemed significantly different. The DEG relationship among different Cas13 variants was shown in Venn network using Evenn and the specific value was shown in upset plot by R package (UpSetR). The *p*-value of venn overlap was calculated by web-based tool DynaVenn with Benjamini-Hochberg adjustment. Gene Ontology (GO) and Kyoto Encyclopedia of Genes and Genomes (KEGG) analysis of genes with absolute FC > 1.5, FDR < 0.05 were performed by R package clusterprofiler with default settings (*p* value < 0.05).

### RIP-seq analysis

The RIP-seq reads were aligned against the hg38 human reference genome using HISAT2. Since the RIP-seq data is strand-specific, the peaks were identified for “+” and “–” strands separately using MACS2, with false-discovery rate (FDR) ≤ 0.01 and arbitrary extension size of 150 bp. Those peaks were regarded as significantly strong peaks when passing the following threshold: pileup > 15 and fold enrichment > 4. To analyze the RIP peak binding profile, the TSS/promoter region was annotated by R package TxDb.Hsapiens.UCSC.hg38.knownGene with upstream/downstream 500bp. The feature distribution for the significant RIP-seq peaks were calculated by annotation function in ChIPseeker package. Motifs for the peak binding regions were calculated using STREME function in MEME-Suite with motif width < 30, *p*-value < 0.05 and shuffled input sequences as background.

### Lasso-based machine learning model design

In one-vector Cas13d screen, only the gRNAs that target non-essential genes were preserved. Among these, partial gRNAs designed for junctions were further filtered. The extended target sequence (± 10 bp on both sides of the spacer region) of the remaining gRNAs were used for the lasso model training. Based on gRNA summary table of one-vector Cas13d screen (Day5 vs. Plasmid), label 1 was set for those gRNAs with LFC < −0.5 and label 0 was set for other gRNAs. Totally, 195 features were generated for each extended target sequence, including one-hot coding for base type in each locus, pair bases content and melting temperature of different fragments. The feature matrix was split into 5 consecutive folds (with shuffling). Each fold was then used once as a validation while the 4 remaining folds form the training set. The Receiver Operating Characteristic (ROC) Curve was drawn and Area Under the ROC Curve (AUC) was calculated for each cross validation. The coefficients (weight vector) were calculated and those base location related weights were used to show the base preference for each locus.

### Statistical analysis

Statistic significances were calculated by GraphPad Prism 8.4.0 and the data were shown as mean ± SD or mean ± SEM. A one-way ANOVA with Dunnett’s multiple comparisons test was used to assess significance between more than two groups. The two-way ANOVA with Tukey’s multiple comparisons test was used to compare differences between groups. Statistical significance was determined by an unpaired two-tailed t-test. Asterisks indicate \**p* < 0.05 and \*\**p* < 0.01.

## ACKNOWLEDGEMENTS

We thank Dr. Tengfei Xiao for technical advice. This work was supported by the National Natural Science Foundation of China (31871344; 32071441), the Fundamental Research Funds for the Central Universities (N182005005; N2020001; N2220001), the 111 Project (B16009), and Liaoning Revitalization Talents Program (XLYC1807212) to T.F.; and the research grant from National Institute of Health (R01HG010753) to W.L.

## AUTHOR CONTRIBUTIONS

T.F. and W.L. conceived the study and designed the research. Zexu.L. and Zihan.L. conducted most of the experiments. Zexu. L. and X.C. conducted bioinformatics analysis. All the authors analyzed the data. T.F. and W.L. wrote the manuscript with the input from Zexu. L., Zihan.L., X.C. and help of all the other authors. T.F. and W.L. supervised the study.

## SUPPLEMENTAL FIGURES

**Figure S1.**
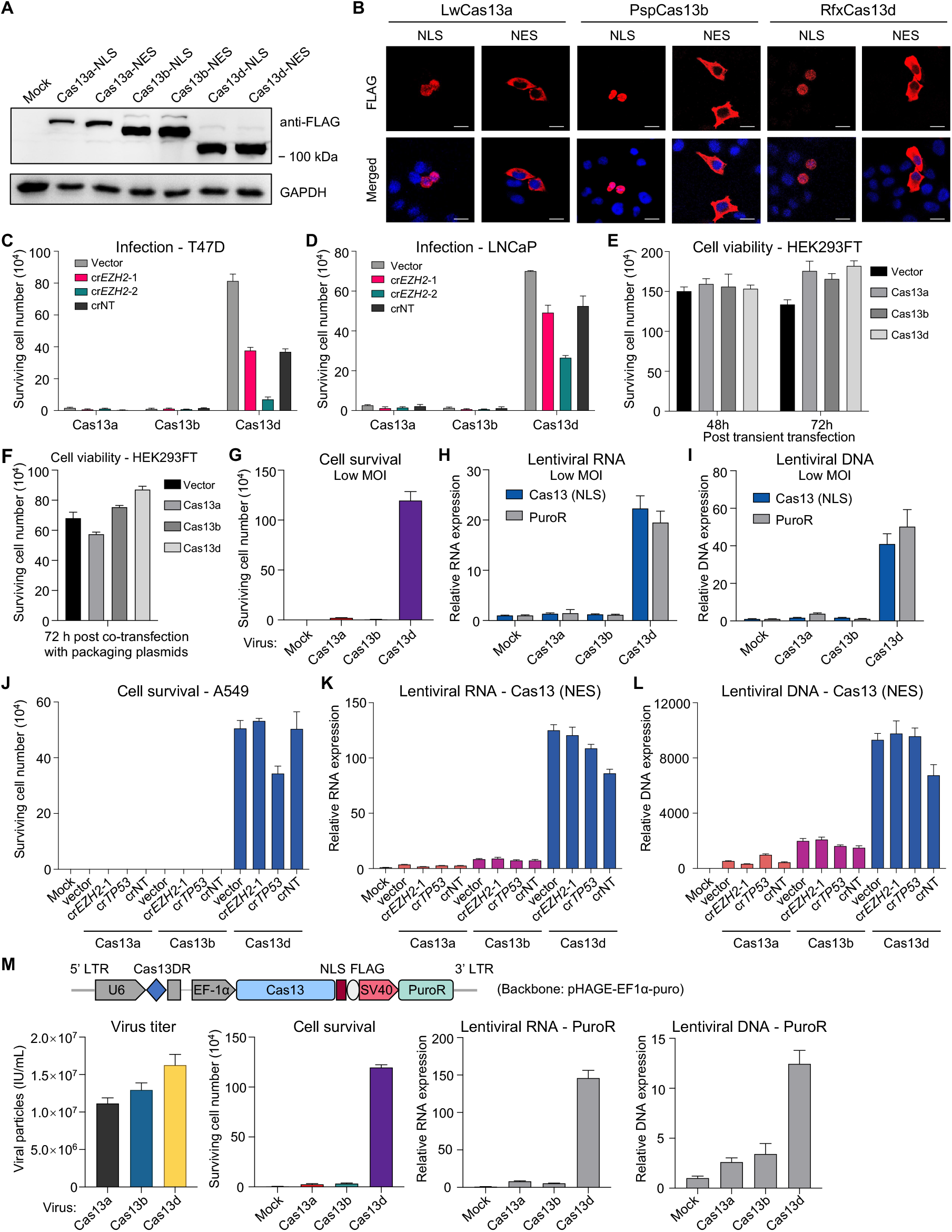
Assessment of lentiviral defect for one-vector Cas13 systems, related to Figure 1. (A) Western blot of protein extracts from HEK293FT cells transiently expressing different Cas13 effectors in one-vector format. (B) Immunocytochemistry of Cas13 proteins showing localization and expression. Scale bar, 20 µm. (C-D) Surviving cell number after puromycin selection for T47D (C) or LNCaP (D) cells infected by indicated one-vector Cas13 lentiviruses, mean ± SEM with n = 3. (E) Cell viability analysis for HEK293FT cells with transient transfection of indicated one-vector lentiviral constructs, mean ± SEM with n = 3. Vector: lentiv2-w/o Cas9. (F) Cell viability analysis for HEK293FT cells during lentivirus production by transient transfection of one-vector Cas13 along with packaging plasmids, mean ± SEM with n = 3. (G-I) Assessment of surviving cell number (G), lentiviral RNA (H) and integrated lentiviral DNA (I) in A549 cells infected with indicated lentiviruses at low MOI. (J-L) Assessment of surviving cell number (J), lentiviral RNA (K) and integrated lentiviral DNA (L) in A549 cells infected with indicated lentiviruses containing different crRNAs. (M) Evaluation of lentiviral defect for one-vector Cas13 systems under pHAGE-EF1α-puro plasmid backbone.

**Figure S2.**
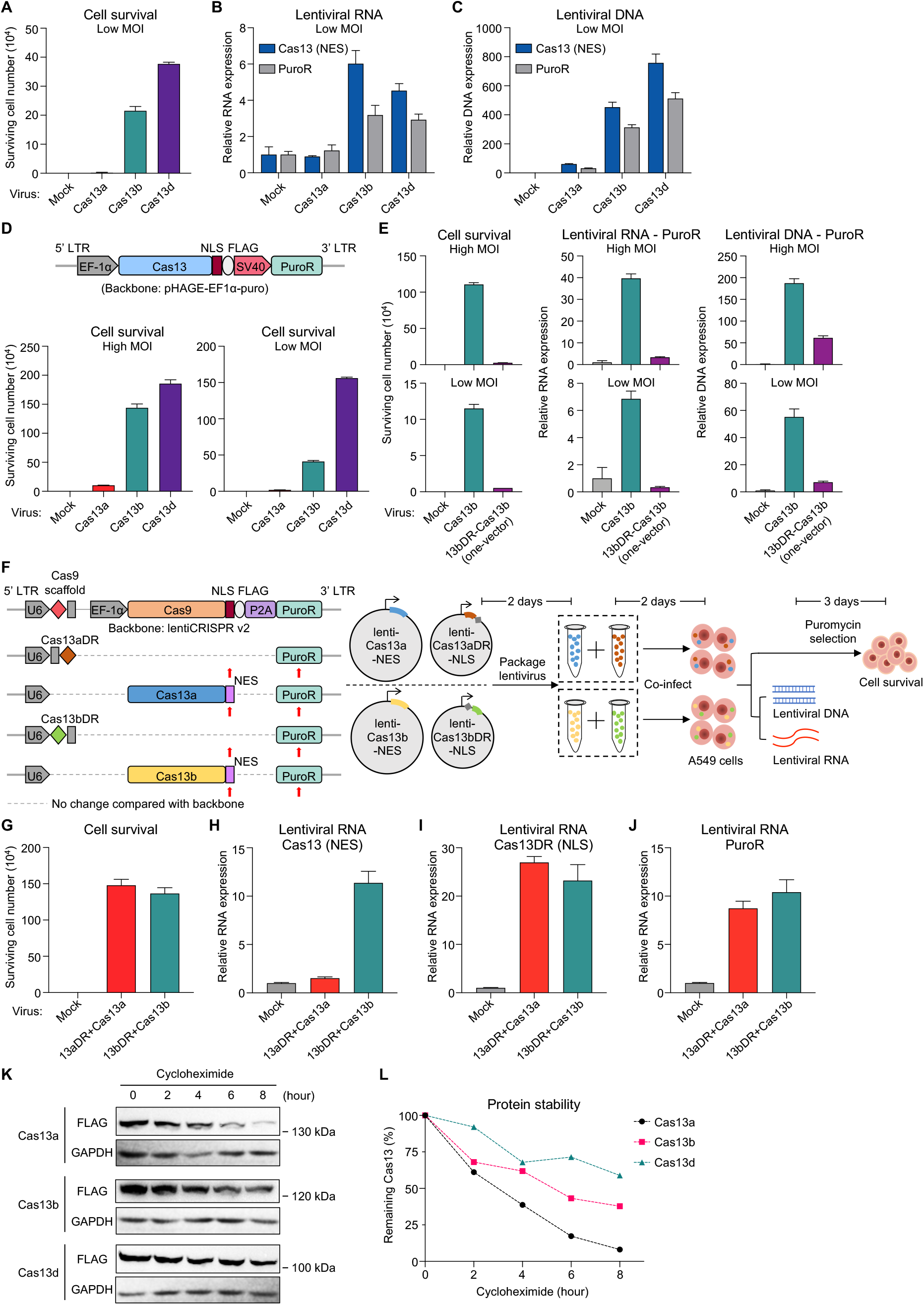
Evaluation of two-vector Cas13 systems for lentiviral defect, related to Figure 2. (A-C) Assessment of surviving cell number (A), lentiviral RNA (B) and integrated lentiviral DNA (C) in A549 cells infected with indicated lentiviruses at low MOI. (D) Effect on cell survival after puromycin selection for A549 cells infected with high or low MOI of Cas13-only lentiviruses of two-vector system under pHAGE-EF1α-puro vector backbone, mean ± SEM with n = 3. (E) Assessment of surviving cell number, lentiviral RNA and integrated lentiviral DNA levels in A549 cells infected with indicated lentiviruses at high or low MOI. (F) Schematic of co-infection assay using indicated lentiviruses to evaluate lentiviral defect. Red arrowheads indicate detection region by qPCR at RNA and DNA levels. (G-J) Assessment of surviving cell number (G) and lentiviral RNA levels using Cas13 (H), Cas13DR (I) or PuroR (J) elements in A549 cells co-infected with indicated lentiviruses. (K-L) Cycloheximide (CHX) chase analysis for determining Cas13 protein stability in HEK293FT cells. Western blot showing the protein levels at indicated time points (K) and quantified band intensity was plotted in (L).

**Figure S3.**
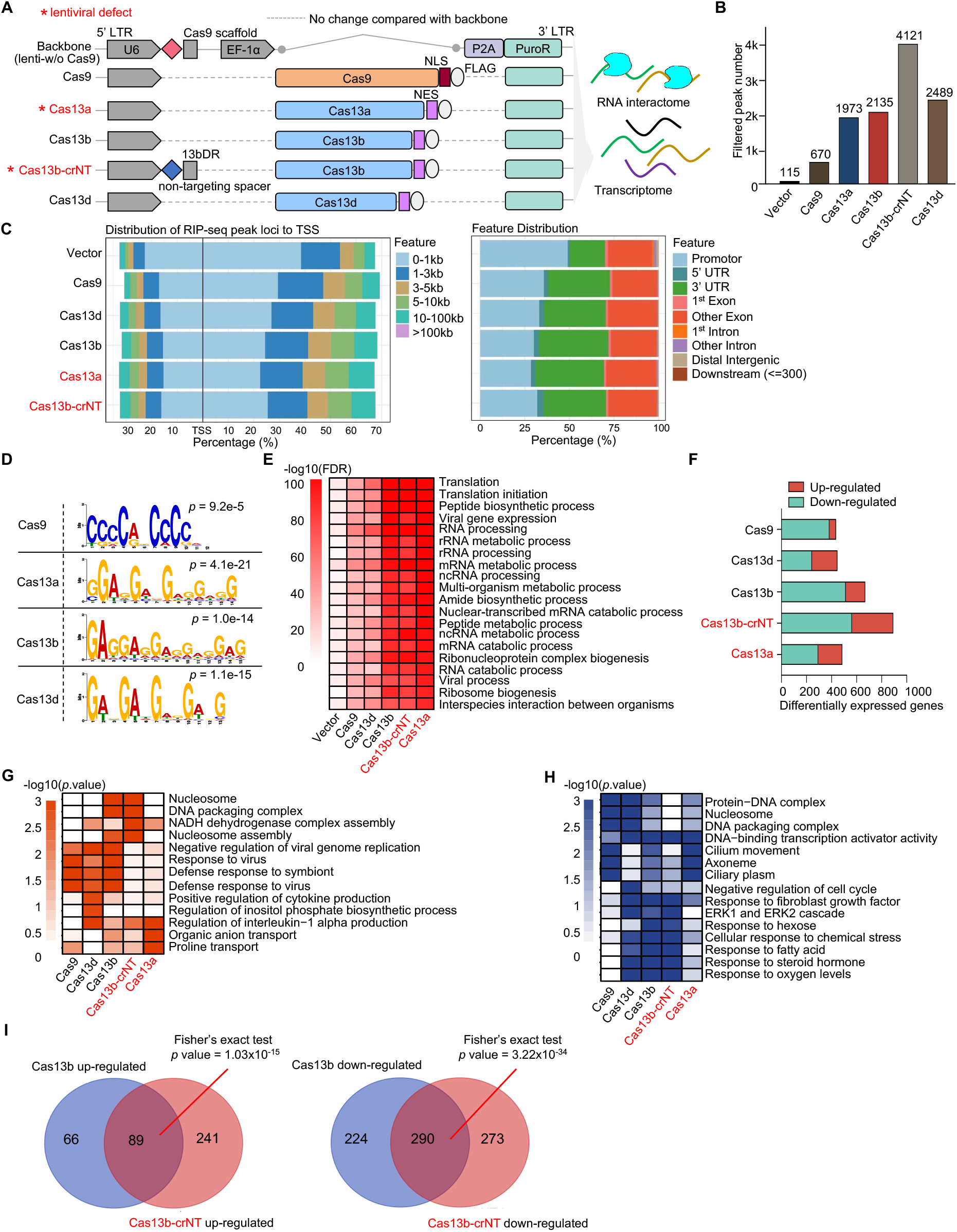
Identification of intrinsic RNA targets of Cas13, related to Figure 3. (A) Schematic of RNA interactome and transcriptome analysis transiently transfected with indicated constructs in HEK293FT cells. Cas13b-crNT indicates a one-vector Cas13b system containing a non-targeting (NT) crRNA. Vector: lentiv2-w/o Cas9. (B) The number of filtered strong RIP binding peaks (pileup > 15; fold enrichment > 4) in different samples. (C) Loci and feature distribution of RIP-seq peaks for indicated samples. (D) Top enriched motifs among RIP-seq peaks for indicated samples. (E) Top GO categories enriched among RIP-seq peak-associated genes across different samples. (F) The number of differentially expressed genes that were up-regulated or down-regulated for indicated samples compared to vector control. (G-H) Top GO categories enriched among up- (G) or down-regulated (H) differentially expressed genes. (I) Venn diagrams representing overlaps of up- (left) or down-regulated (right) genes in Cas13b or Cas13b-crNT groups, versus vector.

**Figure S4.**
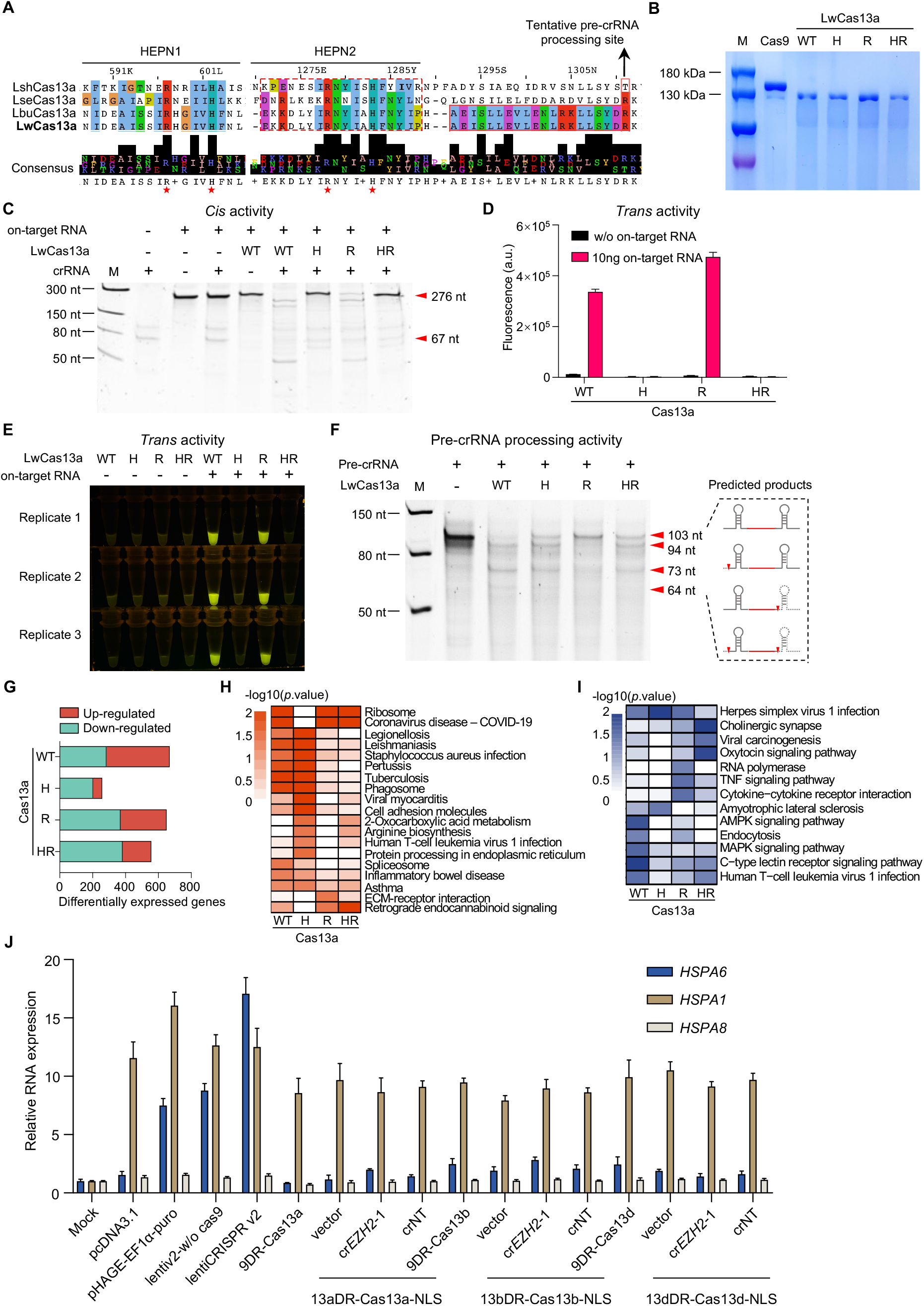
Characterization of Cas13a variants, related to Figure 4. (A) Sequence alignment of catalytic domains for Cas13a derived from different bacteria strains. The conserved functional residues are shown in red star or by black arrow. (B) Coomassie blue staining showing purified Cas9 and Cas13a proteins. (C) *Cis* RNA cleavage by indicated Cas13a variants during *in vitro* assay. Red arrow indicates the band position of on-target RNA or crRNA. (D-E) *Trans* RNA cleavage by indicated Cas13a variants during *in vitro* assay. Results are shown by either fluorescence signal value (D) or direct visualization under blue light illuminator (E). a.u., arbitrary unit. (F) Pre-crRNA cleavage by indicated Cas13a variants during *in vitro* assay. Red arrows indicate band positions of intact and different cleavage patterns of RNA with structural schematic shown in right. (G) The number of differentially expressed genes that were up-regulated or down-regulated among Cas13 variants compared to vector control. (H-I) Top functional categories enriched among up- (H) or down-regulated (I) differentially expressed genes for indicated samples. (J) RNA expression change by qPCR for indicated genes upon transient transfection with indicated constructs.

**Figure S5.**
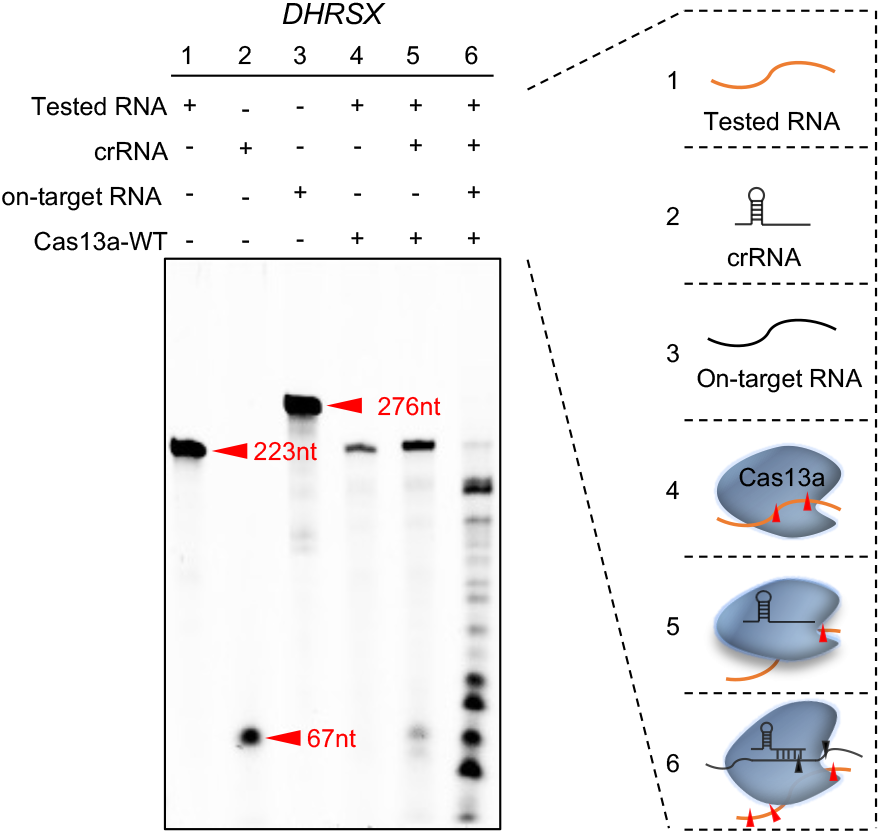
Endogenous RNA targeting by Cas13a, related to Figure 5. *In vitro* assay to assess the effect of Cas13a with differential conformation and complex constitution on endogenous RNA cleavage.

**Figure S6.**
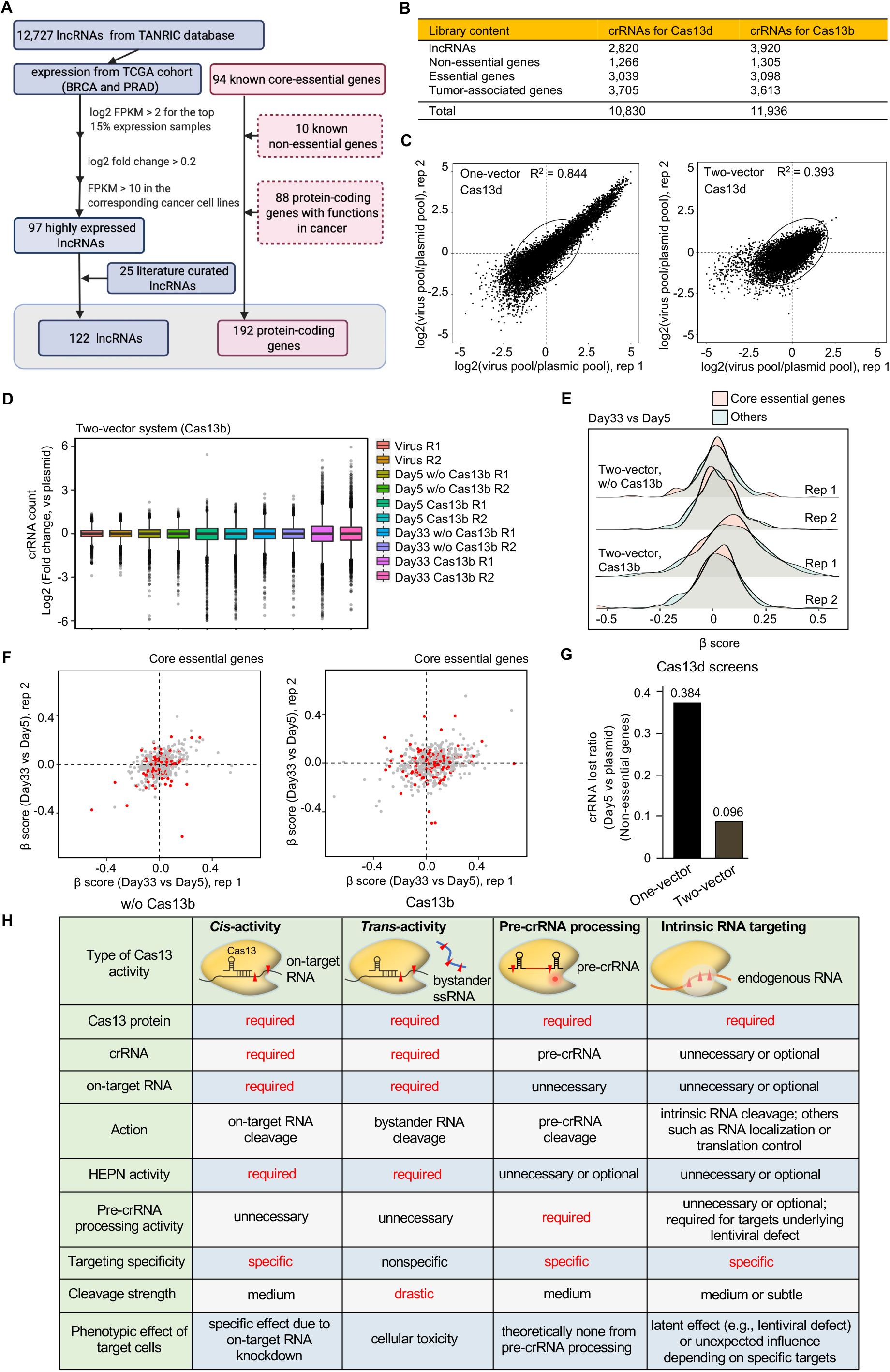
Cas13b/d-based cell fitness screens, related to Figure 6. (A) Schematic of computational pipeline to choose targeted RNA transcripts for Cas13b/d-based cell fitness screens. (B) Statistics of sgRNA libraries for Cas13b/d-based screens. (C-D) Scatter plot showing crRNA abundance change (log2 fold-change) in virus pool versus plasmid library for two biological replicates of either one-vector (C) or two-vector (D) Cas13d screens. R^2^, Pearson’s correlation coefficient. (E) The distribution of crRNA abundance (log2 fold-change) across samples of different time points compared to plasmid pool for two-vector Cas13b screens. (F) Scatter plot of β score change for target transcripts between two biological replicates in Cas13b-null or Cas13b-expressing conditions of two-vector Cas13b screens. Red dots indicate core essential genes. (G) Early loss of crRNAs targeting non-essential genes in one-vector or two-vector Cas13d screens. Log2 fold-change (Day 5 vs. plasmid pool) < −0.5. (H) Comparison of features between different types of Cas13 activities.

## SUPPLEMENTAL TABLES

**Table S1. Oligonucleotides, primers and crRNAs sequences, related to** Figures 1-5**. (attached dataset)**

**Table S2. RIP-seq data, related to** Figures 3 **and** 5**. (attached dataset)**

**Table S3. RNA-seq data, related to** Figures 3, 4 **and** 5**. (attached dataset)**

**Table S4. Cas13 libraries for high-throughput screening, related to** Figure 6**. (attached dataset)**

**Table S5. Cas13 screening data, related to** Figure 6**. (attached dataset)**

